# Brain mitovesicles undergo stage-specific remodelling during the progression of amyloid-β pathology

**DOI:** 10.64898/2026.10.30.762534

**Authors:** Liesbeth Voorbraeck, Jesús Alarcón-Gil, Mite Mijalkov, Massimiliano Passaretti, Zhibai Cao, Doste R. Mamand, Sara Garcia-Ptacek, Julianna Kele, Stephanie L. Fowler, Maria Ankarcrona, Joana B. Pereira, Luana Naia

## Abstract

Mitochondrial dysfunction is an early feature of Alzheimer’s disease that shows nonlinear changes as the disease progresses. However, how these changes are reflected in the mitochondrial material released by brain cells remains poorly understood. Mitovesicles are a recently described extracellular vesicle subpopulation of mitochondrial origin that provide a unique insight into mitochondrial alterations in the brain and could potentially show how nonlinear changes evolve across disease stages of amyloid-β pathology.

Here, we isolated forebrain-derived small extracellular vesicles and mitovesicles from wild-type and *Amyloid precursor protein* (*App)* knock-in (*App^KI^*) mice, carrying familial AD mutations and progressively accumulating amyloid-β plaques, at presymptomatic (2-month) and symptomatic (12-month) stages using high-resolution density gradient fractionation. Vesicle identity and composition were assessed by morphological, biophysical and quantitative proteomic analyses, and mitovesicle respiratory function was measured by extracellular flux analysis.

We show that mitovesicles carry a distinct, predominantly mitochondrial inner membrane proteome that is remodelled by age and disease in a stage-specific manner. Compared with small extracellular vesicles, mitovesicle proteomic profiles captured disease progression more clearly, with symptomatic *App^KI^* mice forming the most molecularly distinct group. At the presymptomatic stage, *App^KI^*mitovesicles showed reduced mitochondrial protein representation, most notably of fatty acid β-oxidation and lipid-metabolism proteins associated with astroglial and vascular signatures, preceding any detectable respiratory impairment. At the symptomatic stage this reversed: *App^KI^*mitovesicles were enriched for oxidative phosphorylation, mitochondrial membrane and transporter proteins, yet showed reduced Complex IV-linked respiration, revealing a dissociation between mitochondrial cargo abundance and respiratory function. Selected cargo changes were not mirrored in whole-brain tissue, indicating selective remodelling of mitovesicle content. Comparison with mitovesicles from post-mortem human Alzheimer’s disease frontotemporal cortex and hippocampus identified a conserved 62-protein mitovesicle core, dominated by oxidative phosphorylation subunits and mitochondrial carriers. In addition, 25 of these proteins were enriched in mitovesicles derived from Alzheimer’s and *App^KI^* forebrains, defining a conserved mitovesicular disease signature.

Together, these findings establish forebrain-derived mitovesicles as a dynamically remodelled vesicle population that reflects progressive mitochondrial dysfunction in the Alzheimer’s disease brain and provides a cross-species resource for future biomarker and mechanistic studies.

## Introduction

Alzheimer’s disease (AD) is the most common neurodegenerative disorder and the leading cause of dementia worldwide. Molecular and metabolic changes in AD start accumulating years before the onset of cognitive symptoms.^1^ Mitochondrial dysfunction is an early metabolic feature of AD, with altered mitochondrial bioenergetics, dynamics and quality control preceding amyloid plaque deposition, tau pathology and neurodegeneration.^2–4^ However, this trajectory is not linear. In *Amyloid precursor protein* (*App)* knock-in (*App^KI^*) mouse models (i.e. *App^NL-F^*and *App^NL-G-F^)*, we showed that mitochondrial hypermetabolism precedes hippocampal amyloid deposition, impaired autophagy and synaptic disorganisation, before transitioning to hypometabolism as pathology advances.^5^ Similarly, mitochondrial bioenergetic signatures distinguish asymptomatic from symptomatic individuals with AD.^4^ Given these stage-dependent alterations, approaches that can resolve mitochondrial alterations at different disease stages are needed.

In recent years, extracellular vesicles (EVs) have emerged as important players in neurodegeneration. EVs are heterogeneous membrane-bound nanovesicles secreted by all cell types. Among them, exosomes and small microvesicles in the 50–150 nm size range are commonly classified as small EVs (sEVs).^6^ Not only have sEVs been shown to mediate the intercellular spread of pathological proteins, including tau filaments^7^ and amyloid β-peptide (Aβ),^8–10^ thereby accelerating disease progression in preclinical models, but they have also attracted considerable interest as a potential source of disease biomarkers.^11,12^ EVs offer two potential advantages over conventional fluid biomarkers. First, the lipid bilayer protects cargo from extracellular proteases and RNases, which may allow more stable and accurate measurement of low-abundance circulating analytes.^13,14^ Second, EV cargo can provide information about the cells of origin and their disease state, with applications in tumour subtyping.^15^ In oncology, plasma EVs have also been used to monitor response to anti-tumour therapies.^16,17^ Beyond diagnostics, their ability to mediate intercellular communication and deliver bioactive cargo has made them attractive candidates as therapeutic vehicles.^6^ Both endogenous^18^ and engineered EVs^19^ can cross the blood-brain barrier to deliver neuroprotective cargo to the brain, making them promising vehicles for the treatment of neurodegenerative diseases.

Recently, we and others have successfully isolated a distinct EV subpopulation enriched for mitochondrial components, which we refer to as mitovesicles.^20,21^ Unlike sEVs, mitovesicles lack the canonical endosomal sorting complexes required for transport (ESCRT) and tetraspanin proteins. They also differ from full mitochondria, with their smaller size, absent cristae and lack of specific mitochondrial proteins such as components of the mitochondrial import machinery (e.g., Translocase of outer mitochondrial membrane 20 (TOMM20)) and Mitofusin 2 (Mfn2)).^21,22^ Importantly, mitovesicles or other EVs containing mitochondria can transfer functional mitochondrial material between brain cells, restoring recipient-cell metabolism and attenuating inflammatory responses in models of neuroinflammation and cerebral ischaemia.^23,24^ Evidence linking mitovesicles to AD is emerging: plasma EVs from patients carry lower levels of mitochondrial Complexes I, III, IV and V,^25^ and mitovesicles from AD brain carry tau filaments, albeit less than sEVs.^7^ Mitovesicles derived from a Down Syndrome mouse model have also been shown to cause synaptic dysfunction.^26^

Although interest in mitovesicles is growing, no study has examined how their proteome and respiratory function change across AD stages, or whether they capture disease-related changes differently from sEVs. Given the stage-dependent mitochondrial trajectory of the *App^KI^* mice, we characterised brain-derived mitovesicles and sEVs from wild-type and *App^KI^* mice at presymptomatic (2-month) and symptomatic (12-month) stages. We found that mitovesicles carry a proteomic signature distinct from sEVs and undergo stage-dependent remodelling that captures disease progression more clearly than the sEV proteome, with a dissociation between mitochondrial cargo and respiratory function at the symptomatic stage. A conserved mitochondrial protein signature was also identified in mitovesicles from post-mortem human AD frontotemporal cortex and hippocampus, supporting the relevance of these findings across species. Taken together, these results suggest that mitovesicle cargo could help unravel the evolving nature of mitochondrial dysfunction by revealing which pathways are affected, while providing a foundation for biomarkers that could complement amyloid and tau markers by capturing a distinct aspect of disease biology.

## Materials and methods

### Animal and ethical considerations

Wild-type (WT) and *App* knock-in mice harbouring Swedish (K670N/M671L), Beyreuther/Iberian (I716F) and Arctic (E693G) mutations (*App^NL-G-F^*hereafter called *App^KI^*), on a C57BL/6J background were studied at 2 and 12 months of age. Sample sizes are specified in figure legends. Procedures complied with Directive 2010/63/EU and were approved by the Regional Ethics Review Board (12779/2019 and 14573-2024); reporting followed ARRIVE guidelines.

### Isolation of sEVs and mitovesicles from mouse brain

sEVs and mitovesicles were isolated from mouse forebrains as previously described by D’Acunzo et al.^27^ Briefly, forebrains were digested, filtered and sequentially centrifuged to finally separate the EV subpopulations by high-resolution iodixanol step-gradient centrifugation. The first three fractions containing microvesicles were discarded, the sEV (fractions 4–6) and mitovesicle-enriched fractions (fractions 7–8) were pooled and centrifuged to wash out residual iodixanol. sEV and mitovesicle pellets were stored at 4°C or -80°C.

### Vesicle size, concentration and morphology

Particle size distributions and concentrations were measured by nanoparticle tracking analysis (NTA) using ZetaView (Particle Metrix). Vesicle morphology and maximum diameter were assessed by cryogenic electron microscopy (cryo-EM) using a Titan Krios microscope (Thermo Fisher Scientific) and Fiji.

### Mass spectrometry (DIA-MS/MS) sample preparation and analysis

Samples were prepared for label-free data-independent acquisition tandem mass spectrometry (DIA-MS/MS) analysis as previously described.^20^ Briefly, peptides were cleared of residual iodixanol by gel purification. Peptides were digested, concentrated, separated by nanoLC and analysed using Orbitrap Exploris with FAIMS Pro™ (ThermoFisher) in DIA mode. Proteomic sample analysis was performed by Biogenity at the Technical University of Denmark. EV fractions included in the analysis were isolated from three male and two female mice per age and genotype. No samples were excluded.

### DIA-MS biostatistics data analysis

Raw DIA-MS data were processed against the *Mus musculus* UniProt database (taxonomy ID: 10090). Downstream analyses were conducted in R (v4.4.2) using Bioconductor 3.20 packages,^28^ including differential enrichment, qualitative differential, co-expression network, overrepresentation and gene set enrichment analysis. To ensure the robustness of the findings, independent research teams applied two different and blinded methodologies to analyse this dataset.

### Human proteomic dataset and cross-species analysis

The MS proteomics data previously published from EV subpopulations of human AD hippocampus and frontotemporal cortex are available via PRIDE repository (PXD037708)^7^. EVs were isolated using the same protocol used for murine vesicles in this study.^27^ Ethical approval, tissue collection and informed consent procedures are described in the original publication. No human tissues were collected or analysed in the present study. The murine and human AD EV datasets were integrated by mapping Human UniProt identifiers to murine orthologs. Consistent with our initial analytical framework, two research teams independently performed different analyses for this cross-species evaluation.

### Western blotting

Proteins isolated from sEVs and mitovesicles (*n* = three/group, female mice) or from total forebrain homogenates (*n* = five/group, 12-month-old female mice) were isolated, loaded on 4–12% Bis-Tris gels and transferred onto nitrocellulose membranes. Odyssey CLx Imaging System (LICORBio) was used to develop the membranes after blocking and incubation with primary and secondary antibodies (**Supplementary table 1**). Image quantification was performed using Fiji.

### Mitovesicle respiratory capacity by Seahorse Analyser

Mitochondrial respiratory capacity of isolated mitovesicles was assessed by measuring oxygen consumption rate (OCR) by Seahorse XF96 analyser (Agilent), following an adapted version of a previously described protocol.^20^

### Statistical analysis

Statistical analysis was done using GraphPad Prism (v10) and data are presented as mean ± SEM. n refers to the number of unique individual biological samples. No blinding nor randomization of samples were performed. Appropriate parametric or non-parametric tests were applied, including unpaired t-tests and one-way or 2way ANOVA followed by Šídák’s multiple comparisons test. Outliers were identified using the ROUT method (Q = 2%). A *P* < 0.05 threshold was considered statistically significant.

Detailed experimental conditions are provided in **Appendix 1**. Reagents and materials are listed in **Supplementary table 2**.

## Results

### Mitovesicles and sEVs differ in morphology but not size, and mitovesicle abundance declines with age

To determine whether the biphasic mitochondrial trajectory of *App^KI^*brains^5^ is reflected in forebrain EVs, we isolated EVs from WT and *App^KI^* mice at 2 and 12 months of age, corresponding to the hypermetabolic presymptomatic and hypometabolic symptomatic stages (**Fig. 1A**).

**Fig. 1.**
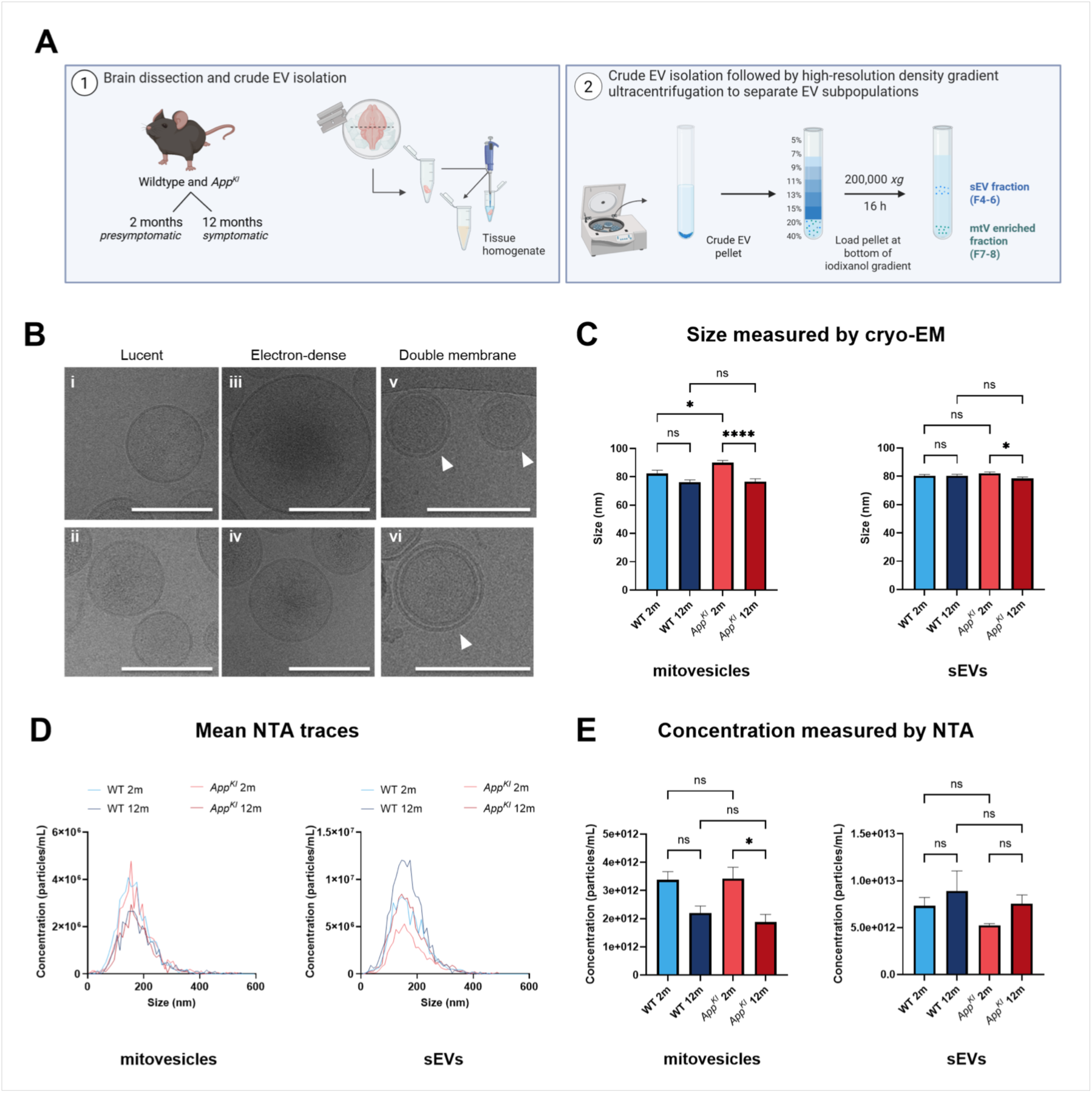
Isolation and structural characterization of forebrain-derived vesicles. (**A**) Schematic representation of the experimental setup. Forebrain-derived vesicles were isolated of 2- and 12-month-old *App^KI^* and wild-type mice by density gradient ultracentrifugation (*n* = 4 per genotype at 2 months, *n* = 3 per genotype at 12 months). Created in BioRender. (**B**) Representative of cryogenic micrographs comparing vesicle morphology; lucent (I–II), electron-dense (III–IV) and double membraned (V–VI). Scale bars = 100 nm (*n* = 61–71 vesicles analysed). (**C**) Quantification of vesicle size measured by cryo-EM separated by vesicle type. (**D**) Mean NTA size-distribution traces (concentration vs. diameter) of each vesicle population. (**E**) Particle concentration of each vesicle population measured by NTA. Data presented as mean ± SEM and statistical significance was done using one-way ANOVA. * *P* ≤ 0.05, **** *P* ≤ 0.0001, ns. non-significant.

Following crude EV isolation, EV subpopulations were separated by high-resolution iodixanol density-gradient ultracentrifugation using a previously established protocol.^27^ sEV- and mitovesicle-enriched fractions (F4–6 and F7–8, respectively) were pooled separately and used for subsequent analyses (**Fig. 1A**).

Following MISEV guidelines,^6^ we performed cryo-EM and NTA, which revealed distinct morphologies between the EV populations: sEVs were predominantly single-membraned and electron-lucent (**Fig. 1B(i**, **ii**)), whereas the mitovesicle-enriched fraction was electron-dense with single- or double-membrane structures (**Fig. 1B(iii–vi**)), the latter showing a ∼6 nm intermembrane space (**Fig. 1B(v**, **vi**)) consistent with the preservation of mitochondrial architecture previously described for intracellular mitochondrial-derived vesicles (MDVs).^21,29^ Despite these structural differences, the two EV populations had comparable overall diameters by cryo-EM, with mean sizes of 80.3 nm for sEVs and 81.3 nm for mitovesicles. Modest age- and genotype-dependent differences were nevertheless detected (**Fig. 1C**); at 2 months, *App^KI^*-derived mitovesicles were larger than WT mitovesicles (*P* = 0.026), whereas this difference was absent at 12 months (*P* > 0.999). Within *App^KI^*mice, both mitovesicles and sEVs were smaller at 12 months than at 2 months (*P* < 0.0001 for mitovesicles; *P* = 0.034 for sEVs; **Fig. 1C**).

By NTA, particle diameter varied with age rather than genotype. sEVs showed an age-related increase in size in WT (142.5 to 159.3 nm; *P* = 0.011) but not *App^KI^* mice (146.8 to 155.9 nm; *P* = 0.231). Mitovesicles showed the opposite genotype pattern, with a significant age-related increase only in *App^KI^*(142.5 to 162.7 nm; *P* = 0.008). Neither EV population differed by genotype at either age (all *P* > 0.800) (**Fig. 1D**, **Supplementary Fig. 1A**). NTA yielded larger particle diameters than cryo-EM (**Fig. 1C**, **Supplementary Fig. 1A**), reflecting the differences between direct measurement of particle dimensions by cryo-EM and estimation of hydrodynamic diameter from Brownian motion by NTA, whose size distributions are influenced by particle heterogeneity and detection sensitivity.^30^

NTA further revealed an age-dependent reduction in mitovesicle concentration in WT (3.38 ± 0.30 to 2.20 ± 0.25 × 10¹² particles/mL; *P* = 0.080) and *App^KI^* mice, where it reached statistical significance (3.43 ± 0.40 to 1.88 ± 0.28 × 10¹² particles/mL; *P* = 0.017). In contrast, sEV concentration did not differ across age or genotype (all *P* > 0.640; **Fig. 1E**).

### Forebrain-derived mitovesicles exhibit a distinct mitochondrial proteomic signature

To understand whether forebrain mitovesicles selectively capture the mitochondrial proteome we profiled the protein cargo of mitovesicle- and sEV-enriched fractions using DIA-MS. Principal component analysis (PCA) revealed a clear separation between sEV and mitovesicle proteomes (**Fig. 2A**). In an additive PERMANOVA model, EV subtype explained the largest share of total proteomic variation (31.4%, *P* = 0.0001), followed by age (5.8%, *P* = 0.0028) and genotype (4.3%, *P* = 0.016). To test whether the genotype effect differed between EV subtypes, we compared *App^KI^*-to-WT centroid distances using a permutation test. At 2 months, distances in mitovesicles and sEVs were similar (mitovesicle-to-sEV ratio = 1.07; *P* = 0.516), whereas at 12 months, *App^KI^*-to-WT centroid distance was 1.41-fold greater in mitovesicle than in sEVs (*P* = 0.045).

**Fig. 2.**
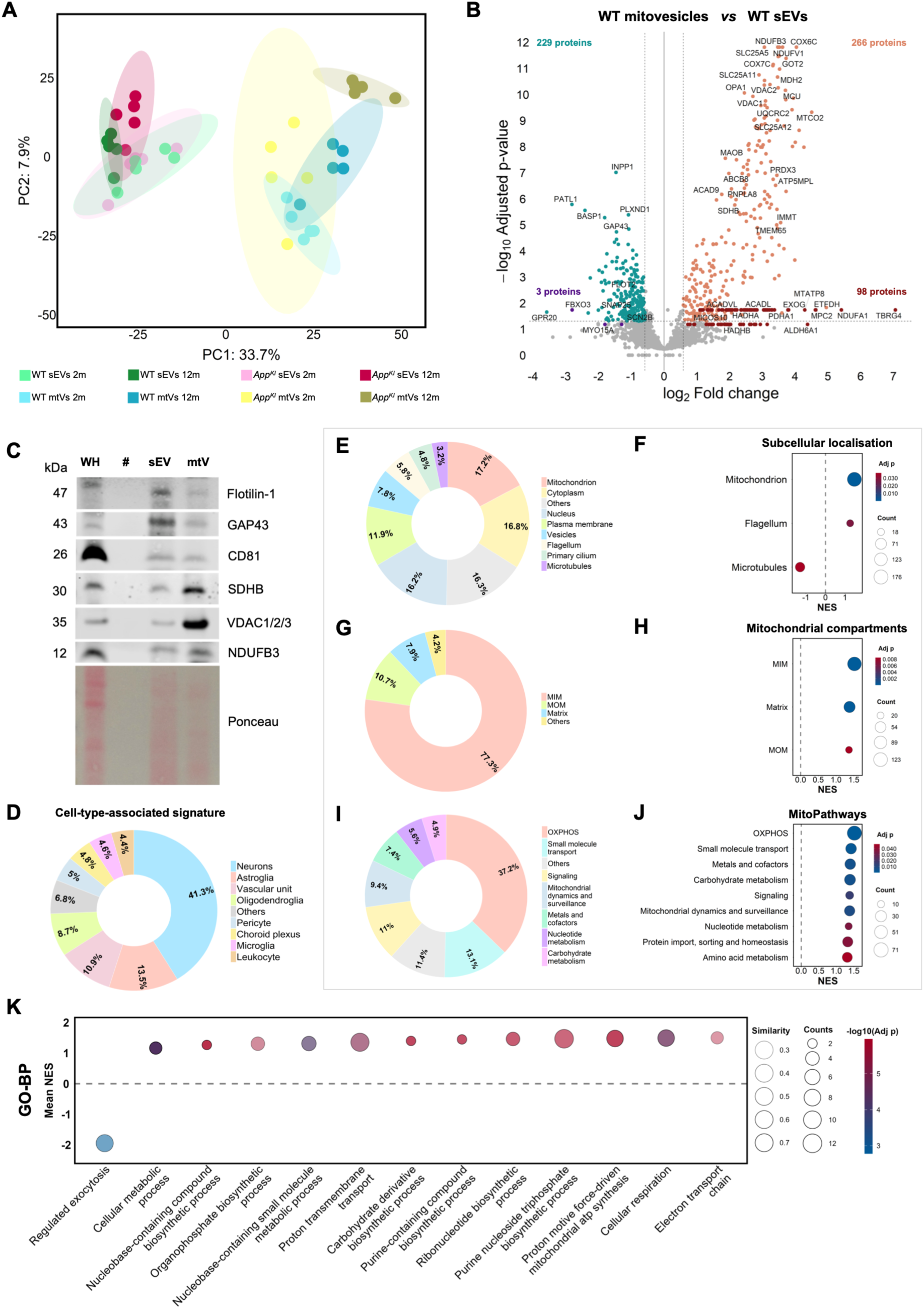
Mitovesicles present with a distinct proteomic signature. (**A**) Principal component analysis (PCA) of the EV proteomic profiles (*n* = 5 per condition). (**B**) Volcano plot depicting differentially expressed proteins between 2-month-old WT mitovesicles and sEVs. The x-axis represents log2 fold change (log2 FC) and the y-axis represents −log10-adjusted *P* values. Blue dots indicate proteins reduced in mitovesicles, purple dots indicate proteins absent from mitovesicles, orange dots indicate proteins enriched in mitovesicles and burgundy dots indicate proteins exclusively detected in mitovesicles. (**C**) Representative western blots of EV-associated (Flot-1, GAP43, CD81) and mitochondria (SDHB, VDAC1/2/3) markers. Ponceau staining was used for normalization and whole-brain homogenate (WH) was included as a positive control. (**D, E, G, I**) Pie charts depicting the distributions of cell-type specific markers (**D**), subcellular organization (**E**), mitochondrial compartments (**G**) and MitoCarta 3.0 MitoPathways (**I**) in the WT mitovesicle fraction at 2 months. (**F, H, J, K**) Dot plots showing enrichment differences between 2-month-old WT mitovesicles and sEVs in subcellular organization (**F**), mitochondrial compartments (**H**), MitoPathways (**J**) and Gene Set Enrichment Analysis (GSEA) analysis of GO-BP terms (**K**). Positive normalized enrichment scores (NES) indicate enrichment in mitovesicles, whereas negative NES indicate enrichment in sEVs. Dot size represents the number of proteins assigned to each term and colour indicates the adjusted *P* value. sEVs: small extracellular vesicles. mtVs: mitovesicles.

To define the molecular features distinguishing mitovesicles from conventional sEVs, we explored the differences between the 2-month-old WT sEV and mitovesicle proteomes. We first compared how often each protein was detected across samples. Three proteins (FBXO3, MYO15A, and ECPAS) were predominantly detected in sEVs but absent from most mitovesicle samples, whereas 98 proteins were primarily detected in mitovesicles but largely undetected in sEVs (**Fig. 2B**, purple and burgundy dots, respectively; *P* < 0.066). Moreover, differential expression analysis revealed 229 proteins reduced and 266 enriched in mitovesicles (**Fig. 2B**, turquoise and orange dots, respectively; *P* < 0.05). This mitovesicular proteomic signature was associated with oxidative phosphorylation (OXPHOS; all five complexes), mitochondrial central dogma, architecture and dynamics, molecule transport and metabolism (e.g., EXOG, OPA1, MICOS10, SLC25A5, PDHA1, MDH2, HADHA/B). Western blotting corroborated some of these findings: GAP43 and flotillin-1 were enriched in sEVs, whereas mitochondrial proteins VDAC1/2/3, NDUFB3 and SDHB were enriched in mitovesicles (**Fig. 2C**).

The 2-month-old WT mitovesicular proteome was predominantly associated with neuronal, astroglial and vascular cell markers, comparable to sEVs (**Fig. 2D**, **Supplementary Fig. 2A**), suggesting that multiple cell populations contribute to forebrain (mitochondrial) vesicle trafficking. Subcellular annotation further confirmed mitochondrial enrichment in mitovesicles (**Fig. 2E** and **F**; *P* < 0.0001), while sEVs contained predominantly proteins assigned to non-mitochondrial cellular compartments and mitochondrial proteins accounting for less than 4% of their proteome (**Supplementary Fig. 2B**). Mitochondrial inner membrane (MIM) proteins accounted for 75.9% of mitochondrial proteins detected in mitovesicles (**Fig. 2G** and **H**; *P* < 0.0001), compared with 31.1% of the complete mouse MitoCarta 3.0 inventory.^31^ This pronounced MIM bias further indicates that mitovesicles do not simply recapitulate the mitochondrial proteome but selectively incorporate mitochondrial cargo associated with the respiratory membrane.

Gene Set Enrichment Analysis (GSEA) converged on OXPHOS, mitochondrial dynamics, small molecule transport and carbohydrate metabolism (*P* < 0.009), among others (*P* < 0.047) (**Fig. 2I-J**). Conversely, synaptic vesicle cycling, exocytosis and actin organization were downregulated in mitovesicles (all *P* ≤ 0.006; **Fig. 2K**, **Supplementary Fig. 2C**), consistent with depletion of presynaptic cargo in the mitovesicle fraction in comparison with sEVs. Gene ontology for cellular components (GO-CC) linked sEV-enriched proteins to synaptic, exocytic and Golgi vesicle membranes, versus respiratory complexes and mitochondrial matrix for mitovesicles (all *P* ≤ 0.047; **Supplementary Fig. 2D**).

Together, these approaches establish forebrain-derived mitovesicles as a distinct EV population selectively incorporating mitochondrial inner-membrane and respiratory-chain cargo. Their greater genotype-associated separation at the symptomatic stage suggests that the mitovesicle proteome captures disease-associated molecular alterations more distinctly than sEVs.

### Unsupervised clustering identifies mitovesicle protein modules associated with lipid metabolism and transmembrane transport

Having established the distinct mitochondrial signature of forebrain-derived mitovesicles, we performed unsupervised hierarchical clustering to understand how the EV proteome is restructured across vesicle subtype, genotype and age. Six protein clusters were defined (**Fig. 3A**). Cluster 6 revealed a conserved mitovesicle core proteome (*n* = 139; **Supplementary Table 3**), in line with the previously observed mitochondrial signature (**Fig. 2**), while the remaining disease-associated clusters showed selective remodelling during amyloidosis.

**Fig. 3.**
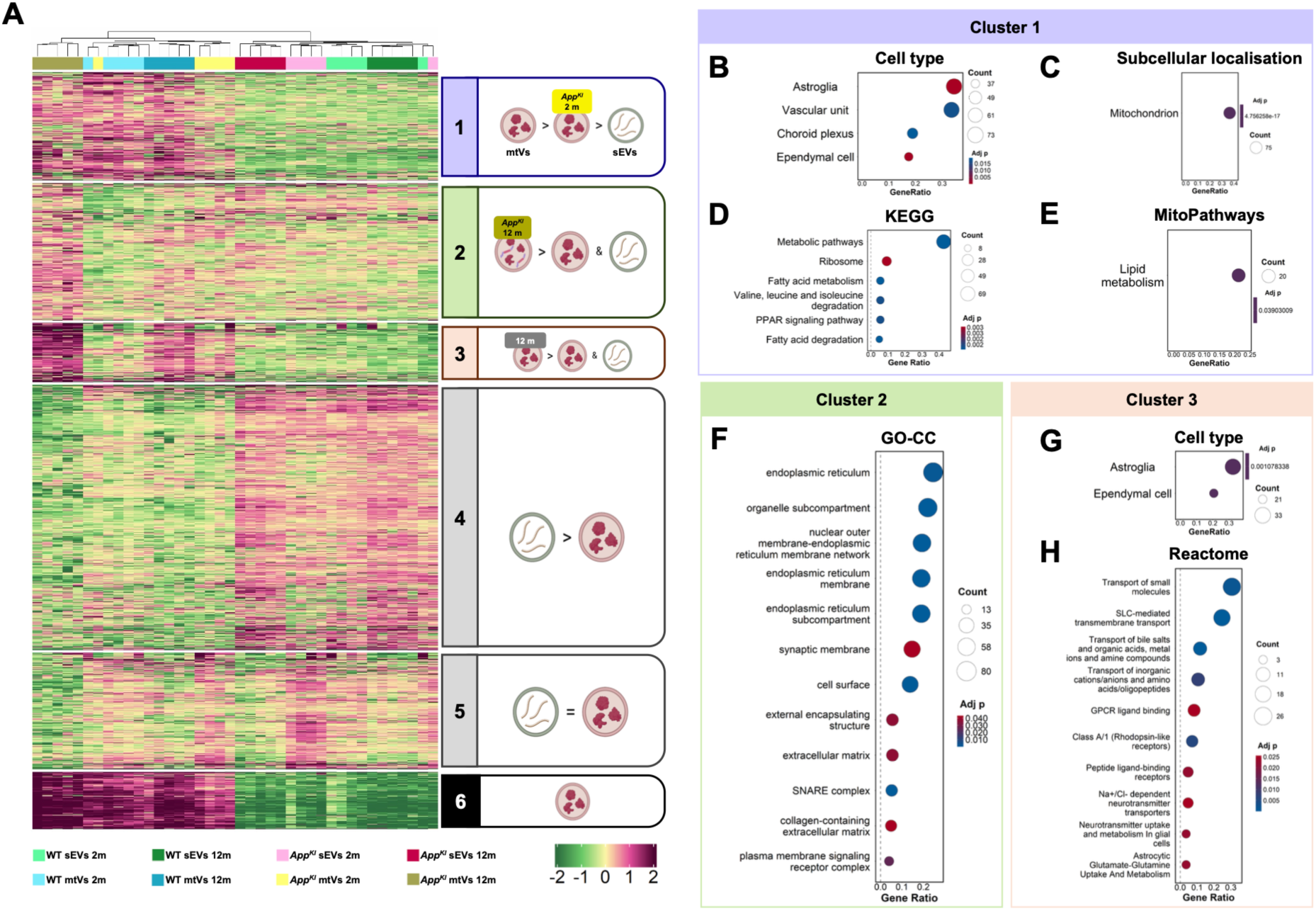
Disease specific mitovesicle proteome profiles emerge during progression of amyloid-β pathology. (**A**) Heatmap of hierarchical clustering of the complete EV proteomic dataset, identifying six protein clusters with distinct abundant profiles. Cluster annotations were created in BioRender. (**B**–**E**) Overrepresentation analysis (ORA) of Cluster 1 showing enrichment by cell-type origin (**B**), subcellular organization (**C**), KEGG pathways (**D**) and MitoPathways (**E**). (**F**) GO-CC ORA analysis of Cluster 2. (**G**, **H**) ORA of Cluster 3, showing enrichment by cell-type origin (**G**) and Reactome pathways (**H**). sEVs: small extracellular vesicles. mtVs: mitovesicles.

Cluster 1 showed overall higher abundance in mitovesicles than in sEVs but lower in presymptomatic *App^KI^* mitovesicles. It was overrepresented for astroglia (*P* < 0.0001), vascular cells (*P* = 0.017), choroid plexus (*P* = 0.019) and ependymal cell (*P* = 0.0002) signatures (**Fig. 3B**), and also for mitochondria (*P* < 0.0001; **Fig. 3C**) and lipid metabolism (*P* = 0.039) annotations, notably fatty acid-related pathways (*P* = 0.002; **Fig. 3D, E**). GO analyses further supported these findings (**Supplementary Fig. 3A** and **B**). Mapping this cluster onto the Reactome pathways ‘Mitochondrial Fatty Acid β-Oxidation’ and ‘Carnitine shuttle’ showed mitovesicles carry most of the catalytic core of the β-oxidation (ACADL, ACADVL, HADHA/B, ACAD11; **Supplementary Fig. 3C**), alongside cardiolipin biosynthetic enzymes (PGS1, PTPMT1) and proteins involved in mitochondrial phospholipid remodelling and lipid metabolism (PNPLA8, FASN, HSD17B10). However, the carnitine shuttle components CPT1A and CPT2 were not detected (**Supplementary Fig. 3D**), suggesting that mitovesicles carry β-oxidation enzymes rather than a complete fatty acid oxidation unit.

Cluster 2 comprised synaptic- and signalling-associated proteins, with a qualitatively increasing trend in 12-month-old *App^KI^* mitovesicles and included endoplasmic reticulum (ER; HSPA5, ITPR1), SNARE complex (STX1A/B, SNAP25, VAMP1) and synaptic membrane (GRIA1, SYP, ADAM22) components (**Fig. 3F**).

Cluster 3 showed higher relative levels in 12-month-old mitovesicles, most strongly in *App^KI^* genotype, and was enriched for astroglial and ependymal signatures (both *P* ≤ 0.002; **Fig. 3G**), including plasma membrane and astrocyte projections (*P* = 0.035 and 0.010, respectively; **Supplementary Fig. 3E**). In addition, this cluster was enriched for small molecule and ion transmembrane transport (*P* ≤ 0.007 and *P* ≤ 0.046, respectively; **Fig 3H**, **Supplementary Fig. 3E**), particularly the mitochondrial SLC carrier family, and for neurotransmitter uptake and metabolism in glia (*P* = 0.025; **Fig. 3H**). To further explore the biological context of this enrichment, we performed a network visualization of this cluster using MitoPathway annotations. Network analysis linked the small molecule transport to OXPHOS in 12-month-old WT mitovesicles, but to detoxification, amino acid metabolism and metals and cofactors pathways in age-matched *App^KI^* mitovesicles, suggesting altered pathway connectivity of transmembrane transport proteins in the context of amyloid pathology (**Supplementary Fig. 3G** and **H**).

Clusters 4 and 5 captured broader vesicle-associated changes: Cluster 4 (sEV-enriched) comprised RNA-binding and heterogeneous nuclear ribonucleoproteins, ER-secretory, ubiquitin-proteasome and redox-related proteins, while Cluster 5 captured a late-stage, vesicle-type-independent reduction in neuronal cytoskeletal, synaptic-trafficking and proteostatic proteins (**Fig. 3A**, **Supplementary Table 3**).

Together, these findings identify early alterations in mitovesicle lipid-metabolism cargo, with an enrichment of astroglial signatures in disease-associated clusters, despite the predominantly neuronal signature of the core mitovesicle proteome, highlighting a previously underexplored dimension of early mitochondrial remodelling during amyloid pathology.

### Co-expression analysis recapitulates stage-dependent remodelling of mitochondrial modules in *App^KI^* mitovesicles

To validate and further dissect these findings, we performed an independent, blinded co-expression network analysis of the forebrain-derived mitovesicle proteome to define protein modules and evaluate how they change across experimental groups. Louvain community detection algorithm applied to the protein-protein correlation matrix identified four distinct co-expression modules, one of which was classified as background noise (**Fig. 4A**).

**Fig. 4.**
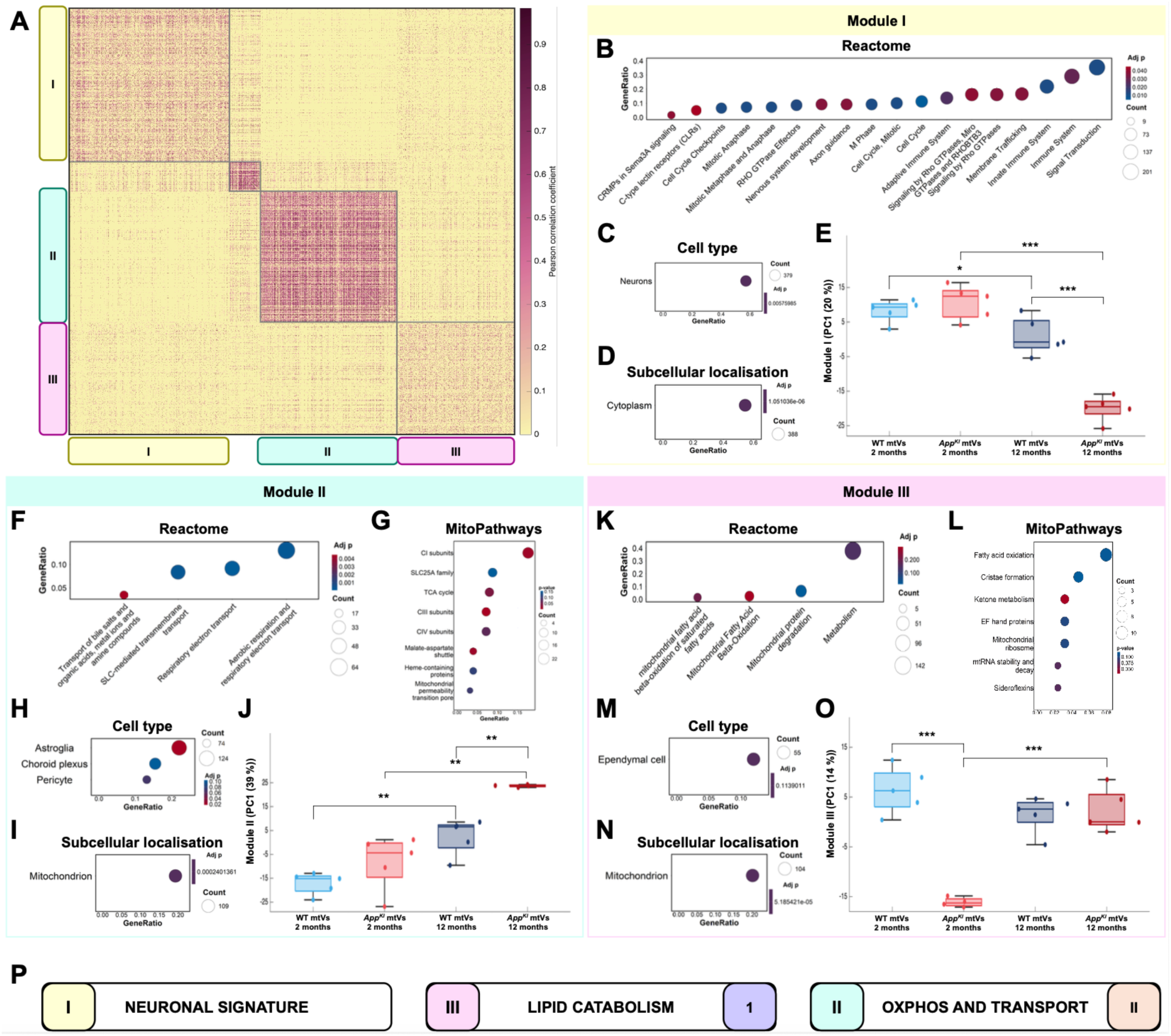
Co-expression modules recapitulate age- and disease-related remodelling in mitovesicles. (**A**) Heatmap of the protein–protein co-expression network. Rows and columns are the 2,348 proteins quantified in all mitovesicle samples (*n* = 20), ordered by module. Colour indicates the Pearson correlation coefficient; only positive correlations ≥ 0.30 are shown and weaker or negative correlations are set to 0 (pale yellow). (**B–D**) ORA of Module I showing enrichment in Reactome pathway (**B**), cell-type origin (**C**) and subcellular localisation (**D**) analyses. (**E**) Boxplot depicting the module eigengene (% variance explained in the axis label) of the different experimental groups for Module I. (**F–I**) ORA of Module II showing enrichment in Reactome pathway (**F**), MitoPathway (**G**), cell-type origin (**H**) and subcellular localisation (**I**) analyses. (**J**) Boxplot showing the module eigengene of the different experimental groups for this module. (**K–N**) ORA of Module III showing enrichment in Reactome pathway (**K**), MitoPathway (**L**), cell-type origin (**M**) and subcellular localisation (**N**) analyses. (**O**) Boxplot showing the module eigengene of the different experimental groups for Module III. (**P**) Schematic showing the relationship between clusters and modules of the two analyses. (E,J,O) Boxes show the median and interquartile range (25th–75th percentile); whiskers extend to the most extreme values within 1.5 × IQR of the box; dots represent individual samples. sEVs: small extracellular vesicles. mtVs: mitovesicles.

Module I was overrepresented in signalling (*P* = 0.011), immune system (*P* = 0.029) and nervous system development (*P* = 0.037), among others (**Fig. 4B**). It was also enriched for neurons and neuronal subcompartments (*P* < 0.0001 and *P* ≤ 0.046, respectively; **Fig. 4C**, **Supplementary Fig. 4A**), and for cytoplasmic and cytoskeletal components (*P* = 0.006 and *P* ≤ 0.046, respectively; **Fig. 4D**, **Supplementary Fig. 4A**), pointing to a neuronal signature. Within mitovesicles, the module’s first principal component declined with age in both genotypes (*P* = 0.044 in WT, *P* < 0.0001 in *App^KI^*) and was further reduced in 12-month-old *App^KI^* mice compared with age-matched WT mice (*P* ≤ 0.0001; **Fig. 4E**).

Module II was enriched for SLC-mediated transmembrane transport (*P* = 0.0003) and respiratory electron transport (*P* < 0.0001; **Fig. 4F**). Subunits of mitochondrial complexes I, III and IV, TCA cycle proteins and members of the SLC25A transporter family tended to be overrepresented (**Fig. 4G**). Module II was also enriched for astroglial signatures (**Fig. 4H**) and mitochondrial (*P* < 0.0001; **Fig. 4I**), plasma membrane (*P* = 0.035) and ER (*P* = 0.031; **Supplementary Fig. 4B**) proteins. The first principal component was higher in 12-month-old compared with 2-month-old mitovesicles in both genotypes, and higher in 12-month-old *App^KI^* than age-matched WT mitovesicles (all *P* = 0.0013; **Fig. 4J**).

Module III was enriched for mitochondrial proteins, including proteins involved in β-oxidation, mitochondrial protein degradation, cristae formation, ketone metabolism and amino acid degradation (**Fig. 4K, L** and **N**, **Supplementary Fig. 4C** and **D**) and for ependymal cell signatures (**Fig. 4M**). Its first principal component was strongly recuced in presymptomatic *App^KI^* mitovesicles compared with presymptomatic WT and symptomatic *App^KI^* mitovesicles (all *P* < 0.0001; **Fig. 4O**).

Together, the two analyses identified co-regulated protein modules within the mitovesicle proteome: a mitochondrial module containing β-oxidation proteins (Cluster 1/Module III), strongly reduced in presymptomatic *App^KI^* mitovesicles, indicating an early genotype effect preceding symptoms, and an OXPHOS and transmembrane transport module (Cluster 3/Module II) that increased with age, most markedly in *App^KI^* mice (**Fig 4P**).

### Dissociation of mitochondrial cargo and respiratory function in symptomatic *App^KI^* mitovesicles

To resolve protein-level alterations across disease progression beyond the module level, we directly compared *App^KI^* and WT mitovesicles at presymptomatic and symptomatic stages. Consistent with the previous clustering and network analyses, genotype-associated differences in mitochondrial cargo were strongly stage-dependent.

At 2 months, *App^KI^* mitovesicles showed reduced mitochondrial representation relative to age-matched WT mitovesicles, affecting both matrix and MIM compartments (**Fig. 5A** and **B**, **Supplementary Fig. 5A** and **B**). This suggests decreased sorting or preferential intracellular clearance at the presymptomatic stage. Differential abundance analysis identified 20 altered proteins: three were increased (CAV1, HPX, BAG5) and 17 decreased in *App^KI^*mitovesicles (**Fig. 5C**), including the mitochondrial fusion protein OPA1, transmembrane transport (SLC25A31, SLC6A6, SLC7A2) and vesicle trafficking (VPS33B, GOLT1B) proteins.

**Fig. 5.**
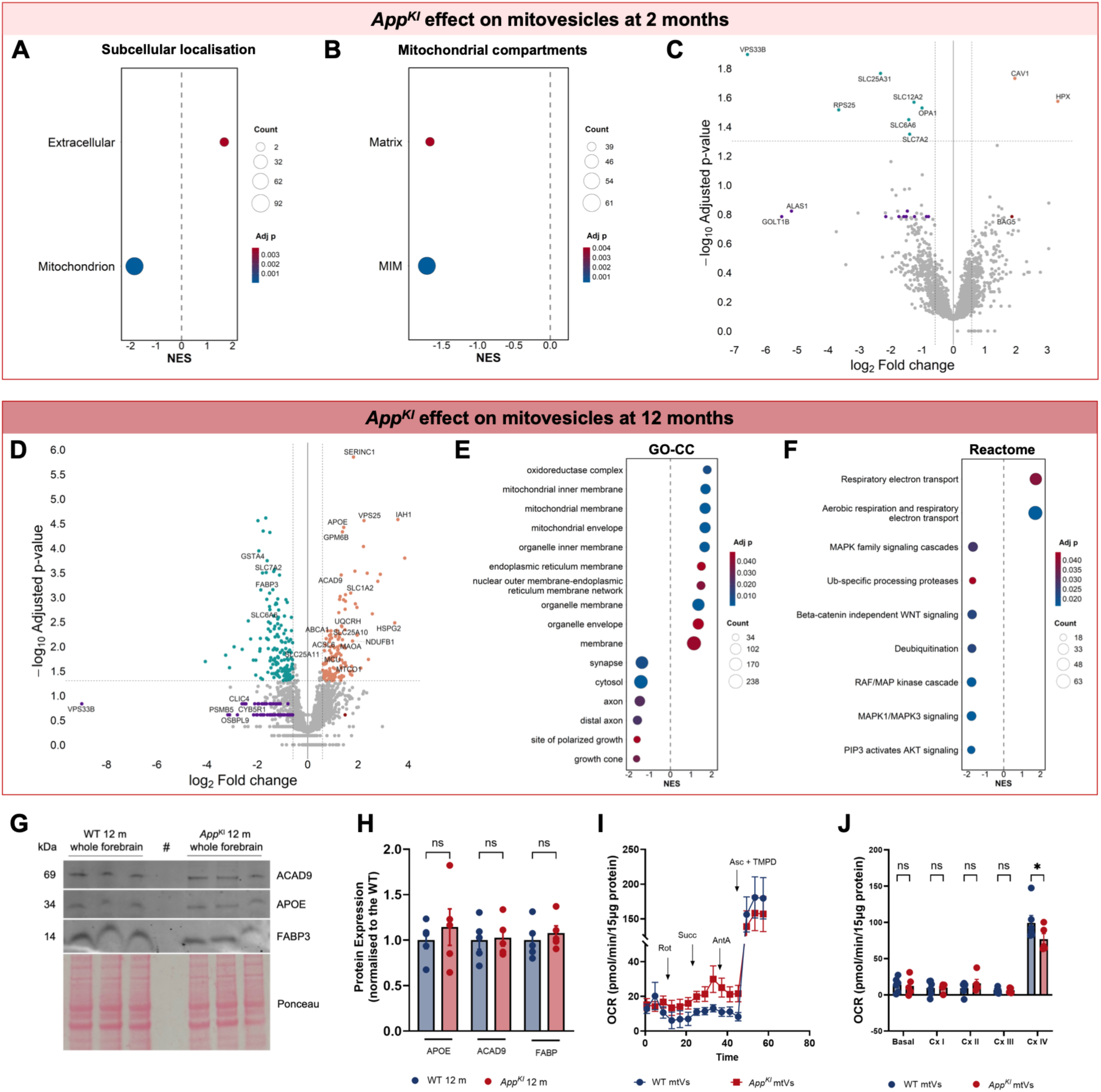
Pairwise analyses reveal age- and disease-dependent changes in mitovesicle proteome. (**A**–**B**) Gene set enrichment analysis of subcellular localization (**A**) and mitochondrial compartments (**B**) in presymptomatic (2 month) WT and *App^KI^* mitovesicles. (**C, D**) Volcano plots showing differentially abundant proteins between WT and *App*^KI^ mitovesicles at the presymptomatic 2-month (**C**) and symptomatic 12-month (**D**) stages. Blue dots indicate proteins significantly reduced in *App*^KI^ mitovesicles, purple dots indicate proteins absent from *App*^KI^ mitovesicles, orange dots indicate proteins significantly enriched in *App*^KI^ mitovesicles and burgundy dots indicate proteins exclusively detected in *App*^KI^ mitovesicles. (**E**–**F**) Dot plots representing GSEA of GO-CC terms (**E**) and Reactome pathway analysis (**F**) in mitovesicles from symptomatic (12 month) WT and *App^KI^*. (**G**) Representative Western blots of selected differentially abundant proteins in whole forebrain homogenates of 12-month-old WT and *App^KI^* mice. (**H**) Quantification of Western blots (*n* = 5 per group). Ponceau staining was used for normalization. (**I**) Seahorse uncoupled oxygen-consumption rate (OCR) traces of mitovesicles from 12-month-old mice following sequential injections of rotenone (2 µM), succinate (10 mM), antimycin A (4 µM) and ascorbate (1 mM)/TMPD (100 mM). Timing of the stimulators/inhibitors indicated on the graph. (**J**) Quantification of basal and respiratory-complex-specific OCR (*n* = 11). Data presented as mean ± SEM and statistical significance was done using one-way ANOVA. * *P* ≤ 0.05, ns. non-significant.

Seahorse extracellular flux analysis confirmed that mitovesicles retain functional respiratory capacity, with measurable but low Complex I–driven basal respiration (**Supplementary Fig. 5C**). Respiration was fuelled with malate and pyruvate, followed by sequential modulation of respiratory chain complexes. At 2 months, basal or Complex I/III/IV–supported respiration did not differ between genotypes (**Supplementary Fig. 5C** and **D**). Succinate elicited no detectable response in either genotype despite the preservation of Complex II subunits (SDHA–D), suggesting that Complex II-supported respiration was not functionally preserved under these assay conditions (**Supplementary Fig. 5D**).

At 12 months, *App^KI^* mitovesicles underwent extensive remodelling: 146 proteins were enriched and 244 reduced relative to age-matched WT mitovesicles (**Fig. 5D**). Genotype-specific presence or absence also increased markedly, with 87 proteins detected exclusively in WT mitovesicles (purple dots in **Fig. 5D**, **Supplementary Fig. 5E**), including membrane trafficking components (CUTA, VPS33B, EXOC5) and proteins involved in β-oxidation (SIK3), lipid metabolism (IMPA1, GPD1L, PGRMC1), amino acid catabolism (GAD2, SLC7A5, GGCT) and carbohydrate metabolism (GSK3A, HAGH, TSC2). Among the proteins detected in both genotypes, ACAD9 and APOE were increased, whereas FABP3 was reduced in symptomatic *App^KI^* mitovesicles (**Fig. 5D**).

Despite this selective loss, GSEA showed an overall increase in mitochondrial representation in symptomatic *App^KI^* mitovesicles, mainly annotated to MIM, mitochondrial envelope, OXPHOS and electron transport pathways and alongside reduced synaptic, axonal and cytosolic compartments and kinase- and ubiquitin-related signalling pathways (all *P* ≤ 0.030; **Fig. 5E** and **F**, **Supplementary Fig. 5F** and **G**). This stage-dependent shift runs opposite to the biphasic metabolic trajectory of *App^KI^*brains,^5^ with less mitochondrial cargo released during the hypermetabolic stage and more during the hypometabolic stage. Thus, despite an overall increase in the representation of mitochondrial proteins, symptomatic *App^KI^*mitovesicles showed selective loss of specific metabolic and vesicle-associated proteins, revealing extensive compositional remodelling.

To assess whether mitovesicle alterations reflected changes in source tissue, we measured ACAD9, APOE and FABP3 in whole-forebrain homogenates. These proteins were selected based on their differential abundance in symptomatic *App^KI^* mitovesicles and their relevance to mitochondrial metabolism, lipid handling and AD pathology.^32–34^ Despite increased ACAD9 and APOE and reduced FABP3 in *App^KI^* mitovesicles, whole-forebrain levels did not differ detectably between genotypes (**Fig. 5G** and **H**), consistent with compartment-specific remodelling of mitovesicle cargo rather than detectable changes in forebrain abundance. Functional analysis further showed that mitovesicles isolated from 12-month-old *App^KI^* forebrain exhibited reduced Complex IV-supported respiration compared with WT mitovesicles (*P* < 0.05; **Fig. 5I** and **J**).

Taken together, symptomatic *App^KI^* mitovesicles carried more mitochondrial protein but showed reduced respiratory capacity, revealing a dissociation between mitochondrial cargo and bioenergetic function.

### Mitochondrial remodelling of mitovesicles tracks amyloid pathology progression rather than physiological ageing

Age-dependent comparisons revealed more extensive remodelling in the disease model, with higher number of differentially expressed proteins (**Supplementary Fig. 5H–N**). Ependymal and astroglial signatures increased at 12 months in both genotypes (*P* ≤ 0.023 in WT; *P* = 0.0002 in *App^KI^*) indicating an age-related shift in cellular representation, whereas the neuronal signature decreased only in symptomatic *App^KI^* mitovesicles (*P* = 0.009; **Supplementary Fig. 5I** and **L**). In WT mitovesicles, ageing was dominated by increased SLC-mediated and small molecule transport with reduced autophagy and receptor tyrosine kinase signalling (all *P* ≤ 0.050) (**Supplementary Fig. 5J**). Age-associated remodelling in *App^KI^* mitovesicles had a mitochondrial-dominated profile with increased MIM and matrix, OXPHOS, carbohydrate and amino acid metabolism representation and reduced synaptic/axonal representation (all *P* ≤ 0.050; **Supplementary Fig. 5M** and **N**), while SLC7A2, SLC6A6 and VPS33B were consistently reduced in *App^KI^* mitovesicles at both ages (**Fig. 5C** and **D**).

Together, these findings suggest that amyloid pathology is associated with mitochondrial remodelling of mitovesicle cargo beyond the changes observed during physiological ageing.

### A mitovesicle core proteome is highly conserved between mouse and human forebrain

To assess whether molecular features of forebrain-derived mitovesicles are conserved across species, we reanalysed a published label-free proteomic dataset of vesicles isolated from post-mortem AD frontotemporal cortex and hippocampus (*n* = 8 for F1–7; *n* = 5 for F8).^7^ Similar to mouse, human mitovesicle proteins were predominantly neuronal (**Fig. 6A** and **2D**), with mitochondria being the most prominent subcellular compartment (**Fig. 6B**) and MIM the predominant compartment (**Fig. 6C** and **2G**). Human mitovesicles were primarily associated with OXPHOS, small molecule transport and signalling pathways, paralleling the composition of murine mitovesicles (**Fig. 6D** and **2I**).

**Fig. 6.**
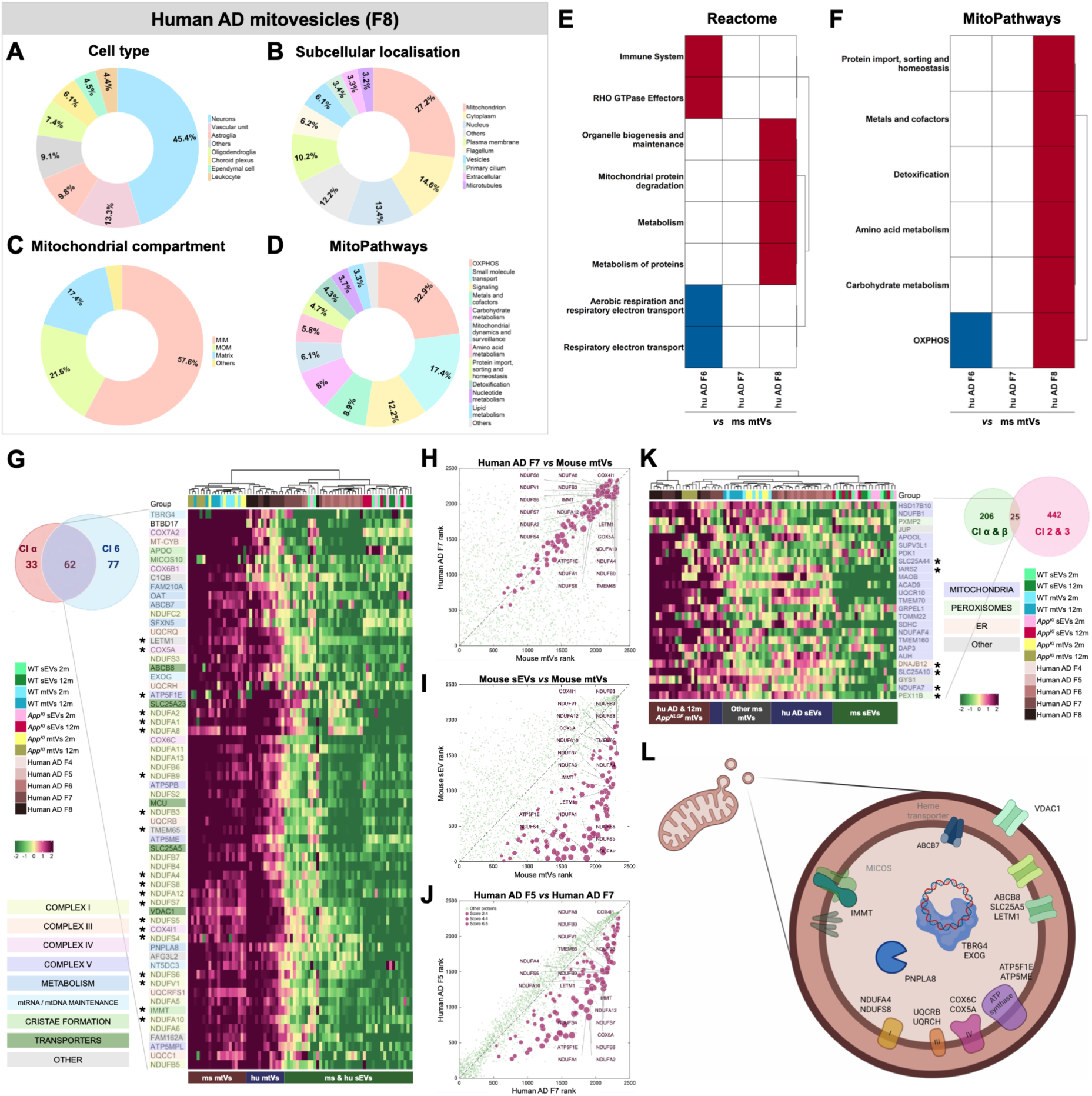
Cross-species comparison identifies a conserved mouse–human mitovesicle proteomic signature. (**A**–**D**) Pie charts showing the inferred cell-type origin (**A**), subcellular distribution (**B**), mitochondrial compartment distribution (**C**) and MitoPathways (**D**) of proteins detected in the human AD mitovesicle-enriched fraction F8 (*n* = 8 for F1–7; *n* = 5 for F8). (**E**, **F**) Comparison of pathway representation in human fractions F6–8 relative to the mouse mitovesicle proteome for Reactome pathway (**E**) and MitoPathway (**F**) analyses. Red indicates higher representation; blue indicates lower representation and white indicates comparable representation. (**G**) Euler diagram showing the overlap between proteins detected in human AD mitovesicle-enriched fractions (Cluster α) and symptomatic *App^KI^* mouse mitovesicles (Cluster 6), after excluding proteins detected in mouse sEVs. The intersection comprises 62 conserved proteins. Hierarchically clustered heatmap showing the abundance profiles of the 62 conserved proteins across the mouse and human mitovesicle fractions. Proteins are colour coded according to selected MitoCarta MitoPathway hierarchy. Stars indicate overlapping proteins in human and mouse mitovesicles. (**H–J**) Rank-based analysis of protein abundance. Protein quantified in all samples were ranked by mean abundance within each group (rank = 1, lowest; rank = 2,348 highest). **(H)** Human fraction F7 versus 12-month-old App^KI^ mouse mitovesicles, showing proteins with concordant ranks across species close to the dashed identity line. **(I)** 12 months App^KI^ mouse sEVs versus mitovesicles, showing proteins ranked higher in mitovesicles than in sEVs below the identity line. **(J)** Human fraction F5 versus F7, showing proteins ranked higher in F7 than in F5 below the identity line. In all plots, magenta dots indicate proteins that met all criteria, which light green dots show all other proteins. The 20 proteins conserved by both approaches are labelled. (**K**) Heatmap showing the overlap of proteins between Clusters 2/3 and Clusters α/β to show the disease-stage related cross-species conserved proteins in human and mouse mitovesicles. Proteins are colour coded according to selected MitoCarta MitoPathway hierarchy. Stars indicate conserved proteins between human and mouse mitovesicles. Euler diagram showing the overlap between proteins detected in human AD mitovesicle-enriched fractions (Cluster α and β) and disease-associated clusters in mouse mitovesicles (Clusters 2 and 3). (**L**) Schematic representation of cross-species conserved mitovesicle cargo. sEVs: small extracellular vesicles. mtVs: mitovesicles.

PCA distinguished sEV- and mitovesicle-associated fractions in both species, with human F7 and F8 largely overlapping, supporting the mouse F7-8 pooling strategy (**Supplementary Fig. 6A**). Comparative enrichment analysis revealed murine mitovesicles and human fraction F7 overlapped substantially in different functional and spatial annotations, while human F8 exhibited greater enrichment in mitochondrial content and some MitoPathway annotations (e.g., amino acid and carbohydrate metabolism; **Fig. 6E** and **F**, **Supplementary Fig. 6B**–**D**).

To explore whether a conserved mitovesicle subproteome is present across both species, we first intersected a cluster of conserved and abundant proteins across human and mouse mitovesicle fractions (Cluster α (95 proteins) in **Supplementary Fig. 6E**) with the previously defined mouse core mitovesicular subproteome (Cluster 6 (139 proteins) in **Fig. 3A**). We identified 62 proteins that were mitovesicle-specific and conserved across species. This mitovesicle core signature was dominated by OXPHOS subunits of Complex I, III, IV and V, mitochondrial solute carriers and proteins involved in cristae formation and metabolism (**Fig. 6G**). Several members of this signature, including VDAC1, have previously been associated with AD,^35,36^ while NDUFS2 and COX5A encoding genes were recently prioritized as candidate AD risk genes through integration of GWAS and brain multi-omics data.^37^ This supports the potential of brain-derived mitovesicle cargo as a source of translational biomarkers of mitochondrial pathology.

To validate this cross-species signature, we independently reanalysed the human and mouse datasets using a rank-based approach. First, we compared protein abundance ranks between mouse mitovesicles and human F7, identifying proteins ranked consistently across species (**Fig. 6H**). Second, to retain only mitovesicle-specific proteins, we compared abundance ranks between mitovesicles and sEV fractions within each species (mouse sEVs and human F5; **Fig. 6I** and **J**). In total, 105 proteins had concordant ranks between mouse mitovesicles and human F7 and were more abundant in mitovesicles than in sEVs in both species. Intersecting these with the 62-protein mitovesicle core proteome identified 20 proteins conserved by both approaches (asterisk in **Fig. 6G**). This high-confidence subset includes several OXPHOS subunits and IMMT, which is downregulated in human AD brains and interacts with the microprotein SHMOOSE (Small Human Mitochondrial ORF Over SErine tRNA).^38,39^ SHMOOSE levels in CSF levels correlate with AD pathology and its genetic variants (SNPs) are linked to higher AD risk.^40^

### Proteins enriched in *App^KI^* mitovesicles are shared with human AD mitovesicles

Lastly, to identify cross-species candidates for an AD-associated mitovesicle signature, we intersected proteins enriched in 12-month-old *App^KI^* mitovesicles (Clusters 2 and 3 in **Fig. 3A**) with proteins detected in both human and mouse mitovesicles (Clusters α and β, the latter comprising proteins most abundant in both symptomatic *App^KI^*and human AD; **Supplementary Fig. 6E**). The 25 AD-related proteins in mitovesicles are mostly of mitochondrial origin and involved in various biological processes, such as OXPHOS (NDUFB1, NDUFAF4, UQCR10, SDHC), TCA cycle (PDK1, SDHC, SLC25A10), lipid (HSD17B10, ACAD9, APOOL) and amino acid metabolism (SLC25A44, MAOB, AUH) and mitochondrial central dogma (SUPV3L1, IARS2, DAP3, HSD17B10) (**Fig. 6K**). Furthermore, six of these candidates were also identified by the previous rank-based analysis of conserved mitovesicular proteins (asterisk in **Fig. 6K**). Several of these proteins have previously been linked to AD, including associations of genetically predicted hippocampal SLC25A10 expression with disease risk and increased IARS2 expression in the blood transcriptome of AD patients.^41–47^ In addition, HSD17B10, ACAD9, GRPEL1, AUH and IARS2 have been detected in CSF from AD patients^48^, supporting their relevance for future studies investigating whether brain mitovesicle signatures can be assessed in biofluids.

Together, our results identify a mitovesicle core proteome conserved between mouse and human forebrain, alongside high-confidence AD-associated cargo candidates for future mechanistic and biomarker studies (**Fig. 6L**).

## Discussion

In this study, we provide a comprehensive characterization of the brain mitovesicle proteome and identify mitovesicles as selectively assembled and dynamically remodelled extracellular carriers that capture changing mitochondrial states during the progression of amyloid-β pathology. Their proteome was strongly enriched for mitochondrial inner-membrane proteins and captured disease-stage-associated changes more distinctly than conventional sEVs. This remodelling followed a biphasic trajectory: At the presymptomatic stage, *App^KI^* mitovesicles showed reduced abundance of β-oxidation and lipid-metabolism proteins associated with astroglial and vascular signatures, whereas symptomatic *App^KI^*mitovesicles accumulated mitochondrial proteins with an enrichment in OXPHOS and small molecule transporter proteins despite reduced respiratory capacity. This late mismatch between mitochondrial cargo abundance and function indicates that greater mitochondrial representation does not reflect enhanced bioenergetic capacity but may instead arise from enhanced sorting of dysfunctional mitochondrial material. We therefore propose that mitovesicle remodelling reflects a transition from early, selective alteration of metabolic cargo to broader disruption of mitochondrial homeostasis as disease progresses. The conservation of a substantial mitochondrial core in human AD brain further supports the cross-species relevance and potential translational value of this extracellular mitochondrial signature.

The selective incorporation of mitochondrial cargo into mitovesicles resembles that of mitochondrial-derived vesicles (MDVs), a mitochondrial quality control system that eliminates mitochondrial components independently of general mitophagy.^29,49^ Proteomic analysis of brain-derived MDVs revealed enrichment of OXPHOS proteins, including respiratory-complex subunits, whereas MDVs generated under oxidative stress contain increased levels of metabolic and redox proteins involved in the TCA cycle, β-oxidation and Fe–S-cluster biology.^50^ These similarities, together with their comparable size and single- or double-membrane architecture, raise the possibility that MDV-mediated cargo selection contributes to the mitochondrial composition of mitovesicles. MDVs delivered to the late endosome or multivesicular bodies could potentially be released extracellularly rather than directed towards lysosomal or peroxisomal degradation, providing one possible route for mitovesicle biogenesis. This route is particularly relevant in AD, in which mitophagy and lysosomal proteolysis are impaired in mouse models and human brain.^51,52^ When intracellular degradative capacity becomes limited, extracellular disposal may provide an alternative route for removing damaged mitochondrial components, potentially accounting for the increased representation of mitochondrial proteins in mitovesicles at symptomatic stages. Once released, mitochondrial components may enter extracellular fluids or be internalized by neighbouring cells for degradation, a process named transmitophagy, which is altered in AD models.^53^

Nevertheless, several MDV-associated proteins are absent from mitovesicle proteomes reported here and by others,^21^ and the two populations should not be considered equivalent. Mitovesicles may represent extracellular descendants of a specific MDV subtype or be a distinct EV population generated through an independent pathway altogether. Because our data define mitovesicle composition but not directly trace their biogenesis, this model should be regarded as an interpretive framework rather than a demonstrated mechanism.

While mitovesicles underwent age-dependent remodelling in both genotypes, their alteration in presymptomatic *App^KI^* may be particularly informative. At 2 months, before detectable plaque deposition or bioenergetic decline, a small but coherent set of proteins involved in β-oxidation, lipid metabolism and cardiolipin remodelling was already reduced in *App^KI^* mitovesicles (Cluster 1/Module III). Rather than reflecting a generalized loss of mitochondrial content, this coordinated change suggests selective decrease of pathways linking lipid availability and mitochondrial membrane integrity to energy production. This early signature is particularly relevant as human genetic studies position lipid processing as a core component of AD susceptibility, alongside Aβ metabolism, tau biology and immunity.^54^

Several altered proteins have also been implicated in AD: FASN, which catalyses *de novo* long-chain fatty acid synthesis, is upregulated in mouse models and patients^55^; HADHA, a component of the mitochondrial trifunctional protein required for long-chain β-oxidation, is increased in AD patients’ cortex but decreased in CSF^48^; and HSD17B10, formerly known as endoplasmic reticulum-associated Aβ-binding protein or ERAB, is a direct Aβ-binding partner and links mitochondrial metabolism to AD-associated dysfunction.^56^ Notably, these proteomic alterations were predominantly enriched in astroglial, vascular, and ependymal signatures, suggesting involvement of neurovascular and brain–CSF interfaces, both implicated in AD-related dysfunction.^57,58^ Astrocytes buffer neuron-derived fatty acids through lipid-droplet storage and mitochondrial and peroxisomal β-oxidation.^59^ Both neuron–astrocyte fatty acid coupling and cerebrovascular function, supported by endothelial cells and pericytes, are compromised early in AD.^57,60,61^ Within this broader context, we previously showed that Aβ-loaded human astrocytes initially preserve ATP by increasing glycolysis and shifting towards lipid droplet-supported, (predominantly) peroxisomal fatty acid oxidation before respiratory capacity declines.^62^ Moreover, we previously showed that astrocyte-derived mitovesicles support neuronal synaptic ATP through mitochondrial β-oxidation,^20^ a pathway affected in presymptomatic *App^KI^*mitovesicles. Together, these findings position lipid-associated mitovesicle remodelling as an early molecular signature of AD brain that may precede bioenergetic loss.

A second trajectory was the progressive enrichment of astroglial- and transporter-associated proteins in mitovesicles, most pronounced in symptomatic *App^KI^* mice. This cluster connected three groups: astrocyte identity and organization proteins – including GFAP, S100B, MAOB and HepaCAM; glutamate and GABA transporters reported to be altered in AD brains (SLC1A2/EAAT2, SLC1A3/EAAT1, SLC6A11/GAT3)^63,64^; and mitochondrial proteins governing substrate exchange and membrane organization. Together, these proteins may reflect reorganization of astrocytic functions, coupling neurotransmitter clearance to metabolic processing, as glutamate and GABA uptake imposes an energetic demand and their metabolism feeds mitochondrial carbon flux through 2-oxoglutarate and the GABA shunt.^65,66^ This interpretation is consistent with single-cell studies showing heterogeneous, stage-dependent astrocyte alterations in neurotransmission, lipid metabolism, antioxidant defence and bioenergetics during AD progression.^67,68^ The transporter-rich mitovesicle phenotype may therefore reflect astrocytic remodelling consistent with an adaptive response to sustained stress. However, increased astrogliosis in symptomatic *App^KI^* mice could also increase the proportion of astroglia-derived mitovesicles independently of cargo remodelling.^5,69^ Thus, the signature may reflect both an altered astrocytic state and a shift in the cellular sources contributing to the mitovesicle pool.

By 12 months, symptomatic *App^KI^* mitovesicles showed broader mitochondrial enrichment, with more OXPHOS-related proteins than presymptomatic *App^KI^* or age-matched WT mitovesicles, alongside reduced vesicle-associated markers. Although mitochondrial dysfunction is a hallmark of healthy ageing,^70^ OXPHOS cargo was less prominent in ageing WT mitovesicles, suggesting that AD phenotype redirects rather than simply accelerates age-dependent remodelling. Notably, increased OXPHOS representation did not indicate preserved function, as symptomatic *App^KI^* mitovesicles displayed reduced Complex IV-supported respiration despite relative preservation of this complex in brain mitochondria in this model.^5^ This function-to-cargo mismatch indicates that increased OXPHOS cargo may arise from increased damaged/oxidised cargo selectivity under mitochondrial stress.^5,71^ Concurrent enrichment of presynaptic/SNARE proteins and ER chaperones further suggests that late-stage mitovesicles capture broader synaptic and proteostatic stress.^72^

The mitovesicle signature extended to human AD forebrain, with 62 proteins conserved across species and independently generated datasets using the same isolation strategy.^7^ These proteins formed a coherent mitochondrial module spanning the respiratory chain (NDUFS2, UQCRFS1, COX5A), cristae organization (IMMT and MICOS10) and metabolite exchange (VDAC1 and SLC25A5). Ultra-deep proteomics in human brain and CSF detected 34 proteins from the conserved core, 13 of which – including several respiratory-chain subunits, OAT, LETM1 and AFG3L – were reduced in AD samples.^48^ This supports the accessibility of individual signature components in CSF. The presence of ependymal- and choroid plexus-associated proteins further suggests a connection with the CSF interface, although CSF entry remains to be demonstrated. Future mitovesicle enrichment and high-plex single-EV approaches^73,74^ could determine whether these proteins coexist within discrete CSF vesicle populations and whether specific mitovesicle subpopulations support disease staging or patient stratification.

Several limitations should be considered. Our findings are primarily compositional and, therefore, future functional studies will be required to establish the consequences of the altered cargo. Moreover, the *App^KI^*model lacks tau pathology and overt neurodegeneration, limiting its representation of human disease.^69^ While density gradient centrifugation enriched mitovesicles, co-isolation of other EV subtypes may contribute to the proteomic profiles, and methodological differences between NTA and cryo-EM limit interpretation of the opposing age-related size trends. The small, AD-only human cohort and incompletely matched brain regions limit the distinction between disease-related and regional effects. Furthermore, independently generated proteomic datasets precluded direct quantitative comparisons across species, so our analyses relied on shared protein identities, abundance ranks and within-species enrichment. The 62-protein core thus represents a conservative set of shared cargo. Finally, larger cohorts will be required to determine whether mitovesicle remodelling differs by sex, particularly given established sex differences in AD risk and progression and growing evidence that lipid metabolism (a prominent axis in our dataset) is differentially affected in women and _men._59,75,76

Together, these findings establish forebrain mitovesicles as a selectively composed and dynamically remodelled EV population that provides an extracellular record of mitochondrial state across ageing and AD progression. Refining their isolation and detection, defining their cellular origins and resolving conserved mitovesicle subpopulations in human biofluids longitudinally will determine whether they can be used to monitor disease trajectories and stratify patients.

## Data availability

The mass spectrometry proteomics data have been deposited in the ProteomeXchange Consortium via the PRIDE partner repository with the dataset identifier PXD085917. The analysis code, original data, processed data, intermediate results and complete results of this study are available at https://github.com/jesusalarcongil/BDMitovesiclesProteomics.

## Acknowledgements

We thank Dr. Takaomi Saido and Dr. Takashi Saito at the RIKEN Center for Brain Science for providing the *App* knock-in mice. We thank the Beta Cell In Vivo Imaging and Extracellular Flux Analysis Core Facility, supported by the Strategic Research Programme in Diabetes, for assistance with the Seahorse experiments. We also thank the Swedish National Cryo-Electron Microscopy Facility, funded by the Knut and Alice Wallenberg Foundation, the Family Erling– Persson Foundation, the Kempe Foundations, SciLifeLab, Stockholm University and Umeå University, for assistance with the cryo-EM experiments. Proteomic sample analysis was performed at the Technical University of Denmark and Biogenity provided the initial proteomic data analysis, including the preparation of figures, tables and reports. We thank Dr. María Latorre Leal at Karolinska Institutet for providing the brain homogenates used in this study. Finally, we thank the late Leif Lundblad for his generous financial support of this work, valuable discussions and longstanding commitment to Alzheimer’s disease research at Karolinska Institutet.

## Supplementary material

Supplementary material is available at *Brain* online.

## References

1. Counts SE, Ikonomovic MD, Mercado N, Vega IE, Mufson EJ. Biomarkers for the Early Detection and Progression of Alzheimer’s Disease. Neurotherapeutics. Jan 2017;14(1):35–53. doi:10.1007/s13311-016-0481-z

2. Terada T, Obi T, Bunai T, et al. In vivo mitochondrial and glycolytic impairments in patients with Alzheimer disease. Neurology. Apr 14 2020;94(15):e1592–e1604. doi:10.1212/WNL.0000000000009249

3. Wang W, Zhao F, Ma X, Perry G, Zhu X. Mitochondria dysfunction in the pathogenesis of Alzheimer’s disease: recent advances. Mol Neurodegener. May 29 2020;15(1):30. doi:10.1186/s13024-020-00376-6

4. Mandal P, Trushina E, Arnold M, Kaddurah-Daouk R, Baloni P. Mitochondrial bioenergetic signatures differentiate asymptomatic from symptomatic Alzheimer’s disease. Commun Biol. Jun 10 2026;doi:10.1038/s42003-026-10474-8

5. Naia L, Shimozawa M, Bereczki E, et al. Mitochondrial hypermetabolism precedes impaired autophagy and synaptic disorganization in App knock-in Alzheimer mouse models. Mol Psychiatry. Sep 2023;28(9):3966–3981. doi:10.1038/s41380-023-02289-4

6. Welsh JA, Goberdhan DCI, O’Driscoll L, et al. Minimal information for studies of extracellular vesicles (MISEV2023): From basic to advanced approaches. J Extracell Vesicles. Feb 2024;13(2):e12404. doi:10.1002/jev2.12404

7. Fowler SL, Behr TS, Turkes E, et al. Tau filaments are tethered within brain extracellular vesicles in Alzheimer’s disease. Nat Neurosci. Jan 2025;28(1):40–48. doi:10.1038/s41593-024-01801-5

8. Perez-Gonzalez R, Kim Y, Miller C, Pacheco-Quinto J, Eckman EA, Levy E. Extracellular vesicles: where the amyloid precursor protein carboxyl-terminal fragments accumulate and amyloid-beta oligomerizes. FASEB J. Sep 2020;34(9):12922–12931. doi:10.1096/fj.202000823R

9. Beretta C, Nikitidou E, Streubel-Gallasch L, Ingelsson M, Sehlin D, Erlandsson A. Extracellular vesicles from amyloid-beta exposed cell cultures induce severe dysfunction in cortical neurons. Sci Rep. Nov 12 2020;10(1):19656. doi:10.1038/s41598-020-72355-2

10. Gabrielli M, Prada I, Joshi P, et al. Microglial large extracellular vesicles propagate early synaptic dysfunction in Alzheimer’s disease. Brain. Aug 27 2022;145(8):2849–2868. doi:10.1093/brain/awac083

11. Taha HB. Alzheimer’s disease and related dementias diagnosis: a biomarkers meta-analysis of general and CNS extracellular vesicles. npj Dementia. 2025/03/12 2025;1(1):3. doi:10.1038/s44400-024-00002-y

12. Huang W, Zhao M, Li L, et al. Extracellular vesicle biomarkers: current status and future perspectives as novel tools in liquid biopsy. Front Immunol. 2026;17:1853671. doi:10.3389/fimmu.2026.1853671

13. Cheng L, Sharples RA, Scicluna BJ, Hill AF. Exosomes provide a protective and enriched source of miRNA for biomarker profiling compared to intracellular and cell-free blood. J Extracell Vesicles. 2014;3doi:10.3402/jev.v3.23743

14. Boukouris S, Mathivanan S. Exosomes in bodily fluids are a highly stable resource of disease biomarkers. Proteomics Clin Appl. Apr 2015;9(3-4):358–67. doi:10.1002/prca.201400114

15. Rontogianni S, Synadaki E, Li B, et al. Proteomic profiling of extracellular vesicles allows for human breast cancer subtyping. Commun Biol. 2019;2:325. doi:10.1038/s42003-019-0570-8

16. Hong CS, Muller L, Whiteside TL, Boyiadzis M. Plasma exosomes as markers of therapeutic response in patients with acute myeloid leukemia. Front Immunol. 2014;5:160. doi:10.3389/fimmu.2014.00160

17. Shao H, Chung J, Balaj L, et al. Protein typing of circulating microvesicles allows real-time monitoring of glioblastoma therapy. Nat Med. Dec 2012;18(12):1835–40. doi:10.1038/nm.2994

18. Yuyama K, Sun H, Sakai S, et al. Decreased amyloid-beta pathologies by intracerebral loading of glycosphingolipid-enriched exosomes in Alzheimer model mice. J Biol Chem. Aug 29 2014;289(35):24488–98. doi:10.1074/jbc.M114.577213

19. Stein JB, Arif M, Cheung H, Lee K-B. Advanced engineering strategies of extracellular vesicles for enhanced central nervous system regeneration and disease modulation. Cell Biomaterials. doi:10.1016/j.celbio.2026.100495

20. Voorbraeck L, Alarcon-Gil J, Giraud R, et al. Astrocyte-neuron mitochondrial transfer via mitoEVs supports neuronal energy metabolism and is impaired in early Alzheimer’s disease. bioRxiv.[Preprint]. doi:10.64898/2026.03.09.710050

21. D’Acunzo P, Perez-Gonzalez R, Kim Y, et al. Mitovesicles are a novel population of extracellular vesicles of mitochondrial origin altered in Down syndrome. Sci Adv. Feb 2021;7(7)doi:10.1126/sciadv.abe5085

22. Sugiura A, McLelland GL, Fon EA, McBride HM. A new pathway for mitochondrial quality control: mitochondrial-derived vesicles. EMBO J. Oct 1 2014;33(19):2142–56. doi:10.15252/embj.201488104

23. Peruzzotti-Jametti L, Bernstock JD, Willis CM, et al. Neural stem cells traffic functional mitochondria via extracellular vesicles. PLoS Biol. Apr 2021;19(4):e3001166. doi:10.1371/journal.pbio.3001166

24. Dave KM, Stolz DB, Venna VR, et al. Mitochondria-containing extracellular vesicles (EV) reduce mouse brain infarct sizes and EV/HSP27 protect ischemic brain endothelial cultures. J Control Release. Feb 2023;354:368–393. doi:10.1016/j.jconrel.2023.01.025

25. Yao PJ, Eren E, Goetzl EJ, Kapogiannis D. Mitochondrial Electron Transport Chain Protein Abnormalities Detected in Plasma Extracellular Vesicles in Alzheimer’s Disease. Biomedicines. Oct 31 2021;9(11)doi:10.3390/biomedicines9111587

26. D’Acunzo P, Argyrousi EK, Ungania JM, et al. Mitovesicles secreted into the extracellular space of brains with mitochondrial dysfunction impair synaptic plasticity. Mol Neurodegener. Apr 14 2024;19(1):34. doi:10.1186/s13024-024-00721-z

27. D’Acunzo P, Kim Y, Ungania JM, Perez-Gonzalez R, Goulbourne CN, Levy E. Isolation of mitochondria-derived mitovesicles and subpopulations of microvesicles and exosomes from brain tissues. Nat Protoc. Nov 2022;17(11):2517–2549. doi:10.1038/s41596-022-00719-1

28. Zhang X, Smits AH, van Tilburg GB, Ovaa H, Huber W, Vermeulen M. Proteome-wide identification of ubiquitin interactions using UbIA-MS. Nat Protoc. Mar 2018;13(3):530–550. doi:10.1038/nprot.2017.147

29. Neuspiel M, Schauss AC, Braschi E, et al. Cargo-selected transport from the mitochondria to peroxisomes is mediated by vesicular carriers. Curr Biol. Jan 22 2008;18(2):102–8. doi:10.1016/j.cub.2007.12.038

30. Vestad B, Llorente A, Neurauter A, et al. Size and concentration analyses of extracellular vesicles by nanoparticle tracking analysis: a variation study. J Extracell Vesicles. 2017;6(1):1344087. doi:10.1080/20013078.2017.1344087

31. Rath S, Sharma R, Gupta R, et al. MitoCarta3.0: an updated mitochondrial proteome now with sub-organelle localization and pathway annotations. Nucleic Acids Res. Jan 8 2021;49(D1):D1541–D1547. doi:10.1093/nar/gkaa1011

32. Nouws J, Nijtmans L, Houten SM, et al. Acyl-CoA dehydrogenase 9 is required for the biogenesis of oxidative phosphorylation complex I. Cell Metab. Sep 8 2010;12(3):283–94. doi:10.1016/j.cmet.2010.08.002

33. Lin YT, Seo J, Gao F, et al. APOE4 Causes Widespread Molecular and Cellular Alterations Associated with Alzheimer’s Disease Phenotypes in Human iPSC-Derived Brain Cell Types. Neuron. Jun 27 2018;98(6):1294. doi:10.1016/j.neuron.2018.06.011

34. Khan W, Aguilar C, Kiddle SJ, et al. A Subset of Cerebrospinal Fluid Proteins from a Multi-Analyte Panel Associated with Brain Atrophy, Disease Classification and Prediction in Alzheimer’s Disease. PLoS One. 2015;10(8):e0134368. doi:10.1371/journal.pone.0134368

35. Manczak M, Reddy PH. Abnormal interaction of VDAC1 with amyloid beta and phosphorylated tau causes mitochondrial dysfunction in Alzheimer’s disease. Hum Mol Genet. Dec 1 2012;21(23):5131–46. doi:10.1093/hmg/dds360

36. Gao H, Jensen K, Nesbitt J, et al. Mitochondrial complex I deficiency induces Alzheimer’s disease-like signatures that are reversible by targeted therapy. Alzheimers Dement. Aug 2025;21(8):e70519. doi:10.1002/alz.70519

37. Lorincz-Comi N, Song W, Chen X, et al. Combining xQTL and genome-wide association studies from diverse populations improves druggable gene discovery. Nature Communications. 2026/02/14 2026;17(1):2801. doi:10.1038/s41467-026-69236-z

38. Adav SS, Park JE, Sze SK. Quantitative profiling brain proteomes revealed mitochondrial dysfunction in Alzheimer’s disease. Mol Brain. Jan 28 2019;12(1):8. doi:10.1186/s13041-019-0430-y

39. Sathe G, Albert M, Darrow J, Saito A, Troncoso J, Pandey A, Moghekar A. Quantitative proteomic analysis of the frontal cortex in Alzheimer’s disease. J Neurochem. Mar 2021;156(6):988–1002. doi:10.1111/jnc.15116

40. Miller B, Kim SJ, Mehta HH, et al. Mitochondrial DNA variation in Alzheimer’s disease reveals a unique microprotein called SHMOOSE. Mol Psychiatry. Apr 2023;28(4):1813–1826. doi:10.1038/s41380-022-01769-3

41. Archer DB, Eissman JM, Mukherjee S, et al. Longitudinal change in memory performance as a strong endophenotype for Alzheimer’s disease. Alzheimers Dement. Feb 2024;20(2):1268–1283. doi:10.1002/alz.13508

42. Tian J, Jia K, Wang T, et al. Hippocampal transcriptome-wide association study and pathway analysis of mitochondrial solute carriers in Alzheimer’s disease. Transl Psychiatry. Jun 10 2024;14(1):250. doi:10.1038/s41398-024-02958-0

43. Chiricosta L, D’Angiolini S, Gugliandolo A, Mazzon E. Artificial Intelligence Predictor for Alzheimer’s Disease Trained on Blood Transcriptome: The Role of Oxidative Stress. Int J Mol Sci. May 7 2022;23(9)doi:10.3390/ijms23095237

44. Han J, Feng GH, Liu HW, Yi JP, Wu JB, Yao XX. Classifying mild cognitive impairment and Alzheimer’s disease by constructing a 14-gene diagnostic model. Am J Transl Res. 2022;14(7):4477–4492.

45. Xi J, He KY, Hou L, et al. Plasma proteomic signatures associate with near-future Alzheimer’s disease dementia conversion in mild cognitive impairment patients. npj Dementia. 2026/01/09 2026;2(1):4. doi:10.1038/s44400-025-00053-9

46. Madrid L, Moreno-Grau S, Ahmad S, et al. Multiomics integrative analysis identifies APOE allele-specific blood biomarkers associated to Alzheimer’s disease etiopathogenesis. Aging (Albany NY). Apr 12 2021;13(7):9277–9329. doi:10.18632/aging.202950

47. Siangphoe U. Bayesian Random-Effects Meta-Analysis Models in Gene Expression Studies. In: Uchiumi F, ed. Gene Expression. IntechOpen; 2022.

48. Wang H, Dey KK, Chen PC, et al. Integrated analysis of ultra-deep proteomes in cortex, cerebrospinal fluid and serum reveals a mitochondrial signature in Alzheimer’s disease. Mol Neurodegener. Jul 25 2020;15(1):43. doi:10.1186/s13024-020-00384-6

49. Soubannier V, McLelland GL, Zunino R, Braschi E, Rippstein P, Fon EA, McBride HM. A vesicular transport pathway shuttles cargo from mitochondria to lysosomes. Curr Biol. Jan 24 2012;22(2):135–41. doi:10.1016/j.cub.2011.11.057

50. Vasam G, Nadeau R, Cadete VJJ, Lavallee-Adam M, Menzies KJ, Burelle Y. Proteomics characterization of mitochondrial-derived vesicles under oxidative stress. FASEB J. Apr 2021;35(4):e21278. doi:10.1096/fj.202002151R

51. Fang EF, Hou Y, Palikaras K, et al. Mitophagy inhibits amyloid-beta and tau pathology and reverses cognitive deficits in models of Alzheimer’s disease. Nat Neurosci. Mar 2019;22(3):401–412. doi:10.1038/s41593-018-0332-9

52. Ye X, Sun X, Starovoytov V, Cai Q. Parkin-mediated mitophagy in mutant hAPP neurons and Alzheimer’s disease patient brains. Hum Mol Genet. May 15 2015;24(10):2938–51. doi:10.1093/hmg/ddv056

53. Lampinen R, Belaya I, Saveleva L, et al. Neuron-astrocyte transmitophagy is altered in Alzheimer’s disease. Neurobiol Dis. Aug 2022;170:105753. doi:10.1016/j.nbd.2022.105753

54. Kunkle BW, Grenier-Boley B, Sims R, et al. Genetic meta-analysis of diagnosed Alzheimer’s disease identifies new risk loci and implicates Abeta, tau, immunity and lipid processing. Nat Genet. Mar 2019;51(3):414–430. doi:10.1038/s41588-019-0358-2

55. Ates G, Goldberg J, Currais A, Maher P. CMS121, a fatty acid synthase inhibitor, protects against excess lipid peroxidation and inflammation and alleviates cognitive loss in a transgenic mouse model of Alzheimer’s disease. Redox Biol. Sep 2020;36:101648. doi:10.1016/j.redox.2020.101648

56. Houfkova A, Schmidt M, Benek O, et al. New insights into the 17beta-hydroxysteroid dehydrogenase type 10 and amyloid-beta 42 derived cytotoxicity relevant to Alzheimer’s disease. Alzheimers Res Ther. Jul 23 2025;17(1):170. doi:10.1186/s13195-025-01821-8

57. Sun N, Akay LA, Murdock MH, et al. Single-nucleus multiregion transcriptomic analysis of brain vasculature in Alzheimer’s disease. Nat Neurosci. Jun 2023;26(6):970–982. doi:10.1038/s41593-023-01334-3

58. Tijms BM, Vromen EM, Mjaavatten O, et al. Cerebrospinal fluid proteomics in patients with Alzheimer’s disease reveals five molecular subtypes with distinct genetic risk profiles. Nat Aging. Jan 2024;4(1):33–47. doi:10.1038/s43587-023-00550-7

59. Maioli S, Nalvarte I, Ankarcrona M, et al. Bioenergetics and lipid metabolism in Alzheimer’s disease: From cell biology to systemic health. J Intern Med. Jan 2026;299(1):20–43. doi:10.1111/joim.70036

60. Qi G, Mi Y, Shi X, Gu H, Brinton RD, Yin F. ApoE4 Impairs Neuron-Astrocyte Coupling of Fatty Acid Metabolism. Cell Rep. Jan 5 2021;34(1):108572. doi:10.1016/j.celrep.2020.108572

61. Du Y, Sun C, Wu L, et al. Oral GLP-1 receptor agonist promotes astrocyte-neuron lactate and lipid transfer with neuroprotective effects. Cell Metab. Aug 4 2026;38(8):1633–1649 e7. doi:10.1016/j.cmet.2026.05.014

62. Zysk M, Beretta C, Naia L, et al. Amyloid-beta accumulation in human astrocytes induces mitochondrial disruption and changed energy metabolism. J Neuroinflammation. Feb 20 2023;20(1):43. doi:10.1186/s12974-023-02722-z

63. Scott HA, Gebhardt FM, Mitrovic AD, Vandenberg RJ, Dodd PR. Glutamate transporter variants reduce glutamate uptake in Alzheimer’s disease. Neurobiol Aging. Mar 2011;32(3):553 e1-11. doi:10.1016/j.neurobiolaging.2010.03.008

64. Wood OWG, Yeung JHY, Palpagama TH, Turner C, Waldvogel HJ, Faull RLM, Kwakowsky A. Upregulated excitatory amino acid transporter 1 (EAAT1) expression in the human medial temporal lobe in Alzheimer’s disease. Neuroscience. Feb 6 2025;566:87–96. doi:10.1016/j.neuroscience.2024.12.027

65. Andersen JV, Schousboe A. Milestone Review: Metabolic dynamics of glutamate and GABA mediated neurotransmission - The essential roles of astrocytes. J Neurochem. Jul 2023;166(2):109–137. doi:10.1111/jnc.15811

66. Pellerin L, Magistretti PJ. Glutamate uptake into astrocytes stimulates aerobic glycolysis: a mechanism coupling neuronal activity to glucose utilization. Proc Natl Acad Sci U S A. Oct 25 1994;91(22):10625–9. doi:10.1073/pnas.91.22.10625

67. Serrano-Pozo A, Li H, Li Z, et al. Astrocyte transcriptomic changes along the spatiotemporal progression of Alzheimer’s disease. Nat Neurosci. Dec 2024;27(12):2384–2400. doi:10.1038/s41593-024-01791-4

68. Sadick JS, O’Dea MR, Hasel P, Dykstra T, Faustin A, Liddelow SA. Astrocytes and oligodendrocytes undergo subtype-specific transcriptional changes in Alzheimer’s disease. Neuron. Jun 1 2022;110(11):1788–1805 e10. doi:10.1016/j.neuron.2022.03.008

69. Saito T, Matsuba Y, Mihira N, et al. Single App knock-in mouse models of Alzheimer’s disease. Nat Neurosci. May 2014;17(5):661–3. doi:10.1038/nn.3697

70. Lopez-Otin C, Blasco MA, Partridge L, Serrano M, Kroemer G. The hallmarks of aging. Cell. Jun 6 2013;153(6):1194–217. doi:10.1016/j.cell.2013.05.039

71. Du F, Yu Q, Yan S, et al. PINK1 signalling rescues amyloid pathology and mitochondrial dysfunction in Alzheimer’s disease. Brain. Dec 1 2017;140(12):3233–3251. doi:10.1093/brain/awx258

72. Anschuetz A, Schwab K, Harrington CR, Wischik CM, Riedel G. A Meta-Analysis on Presynaptic Changes in Alzheimer’s Disease. J Alzheimers Dis. 2024;97(1):145–162. doi:10.3233/JAD-231034

73. Wu D, Yan J, Shen X, et al. Profiling surface proteins on individual exosomes using a proximity barcoding assay. Nat Commun. Aug 26 2019;10(1):3854. doi:10.1038/s41467-019-11486-1

74. Wu B, Yang X, Cai Y, et al. Proteomic profiling of single extracellular vesicles as a promising new approach for the diagnosis and treatment modality of advanced ovarian cancer. NPJ Precis Oncol. Jan 21 2026;10(1):65. doi:10.1038/s41698-026-01271-x

75. Ferretti MT, Iulita MF, Cavedo E, et al. Sex differences in Alzheimer disease - the gateway to precision medicine. Nat Rev Neurol. Aug 2018;14(8):457–469. doi:10.1038/s41582-018-0032-9

76. Arnold M, Nho K, Kueider-Paisley A, et al. Sex and APOE epsilon4 genotype modify the Alzheimer’s disease serum metabolome. Nat Commun. Mar 2 2020;11(1):1148. doi:10.1038/s41467-020-14959-w

