## Supplementary data for "Brain mitovesicles undergo stage-specific remodelling during the progression of amyloid-β pathology"

**Supplementary material**

—

### Extended materials and methods

#### Animal and ethical considerations

Wild-type (WT) and *App* knock-in mice carrying Swedish, Beyreuther/Iberian, and Arctic mutations (*App*$^{NL\text{-}G\text{-}F}$ hereafter called *App*$^{KI}$), with the same C57BL/6J genetic background were used. Both male and female mice at 2- and 12-months-of-age were included. Animals were bred and housed at the Comparative Medicine's Annex (KM-A) at Karolinska Institutet under conditions of controlled temperature (22–23°C), in 12:12 h light/dark cycles and specific-pathogen-free conditions. Cages were enriched in corn-husk nesting material and paper rolls. Food and water were available *ad libitum*. All experimental procedures were carried out in accordance with the guidelines of the Institutional Animal Care and Use of Committee, the European Community directive (2010/63/EU) and protocols approved by the Regional Ethics Review Board (authorization no. 12779/2019 and 14573-2024) in line with the ARRIVE guidelines. All efforts were made to minimize animal suffering and reduce the number of animals used. Mice were sacrificed by cervical dislocation under anaesthesia with 5% isoflurane, brains were collected and kept at -80°C or at 4°C in hibernate-A medium.

#### Isolation of sEVs and mitovesicles from mouse brain

Isolation of brain sEVs and mitovesicles was performed using a high-resolution iodixanol step-gradient ultracentrifugation as previously described by D'Acunzo et al.[1] Briefly, forebrains were treated for 15 min with 18 units/mL papain in Hibernate-A at

37°C. The dissociation was stopped using 1 mM PMSF with protease and phosphatase inhibitors (1:10,000 dilution). The digested brain tissue was centrifuged at 300 *xg* for 10 min at 4°C and the supernatant was sequentially filtered through a 40 μm mesh filter and 0.22 μm syringe filters. The filtrate was subjected to a sequential centrifugation at 4°C at 2,000 ×*g* for 10 min, 10,000 ×*g* for 30 min and at 100,000 ×*g* for 70 min. The EV pellet was resuspended in 35 mL PBS and centrifuged again at 100,000 ×*g* for 70 min at 4°C. EV subpopulations were separated by high-resolution iodixanol (OptiPrep™) step-gradient centrifugation at 200,000 ×*g* for 16 h, at 4°C (SW40Ti Swinging-Bucket rotor, Beckman Coulter). The first three fractions containing microvesicles were discarded, while the sEV (fractions 4–6) and mitovesicle-enriched fractions (fractions 7–8) were pooled, resuspended in PBS and centrifuged two to three times at 100,000 ×*g* for 70 min to wash out residual iodixanol (SW40Ti Swinging-Bucket rotor and TLA-110 Fixed-Angle rotor, Beckman Coulter). sEV and mitovesicle pellets were stored at either 4°C or -80°C.

#### Nanoparticle tracking analysis (NTA)

NTA was performed to determine the particle size distribution and concentration of sEV and mitovesicle fractions using the Particle Metrix ZetaView system, following previously described protocols. Prior to analysis, samples were diluted in PBS filtered through a 0.22 μm membrane to a final volume of 1 mL. Optimal sample dilutions were determined through preliminary measurements to achieve approximately 140–200 particles per frame. Measurements were conducted using the manufacturer's recommended settings for extracellular vesicle analysis. Each sample was analysed in two measurement cycles across 11 cell positions, with 60 frames captured per position using the high-video setting. The acquisition parameters included enabled autofocus, a camera sensitivity of 80, a shutter speed of 100, a scattering intensity of 4.0, and a maintained cell temperature of 24°C. Following acquisition, the recorded data were processed and analysed using ZetaView software (version 8.06.01).

#### Cryogenic electron microscopy (cryo-EM)

Cryo-EM was used for the structural analysis of the vesicles. R-23Cu mesh Quantifoil grids were glow-discharged (40 mA, 60 s) in a GloQube (Quorum). After isolation, the vesicles were resuspended in 30 μL EV-depleted Tris-HCl buffer (pH 7.2). Three μL of the EV solution was applied to the grid, blotted with filter paper (3 s) and then plunge-frozen in liquid ethane using a Vitrobot™ Mark IV (ThermoFisher). Images were obtained with Titan Krios (ThermoFisher) equipped with a K3 direct electron detector (DED) (Gatan) at 64,000× magnification. Data were acquired with a total accumulated electron dose of ~55 $e^{-}/Å^2$ and a nominal defocus range of −3 to −4 µm. Micrographs were analysed using Fiji. Vesicle size was determined by the maximum diameter of each vesicle.

#### Mass spectrometry (MS) sample preparation and analysis

Samples were prepared for MS analysis as previously described.[2] Briefly, residual iodixanol was cleared from the purified fractions by running them on a 12% Bis-Tris gel into a single 1-cm band. Proteins were visualized using EZBlue™. Excised bands were destained (50% acetonitrile/50 mM ammonium bicarbonate), dehydrated (100% acetonitrile), reduced and alkylated (10 mM TCEP, 40 mM CAA, 80 mM ammonium bicarbonate, 30 min) and dehydrated again before overnight trypsin digestion (25 ng/µL) at 37°C. Peptides were extracted (5% formic acid, 50% acetonitrile), concentrated in an SpeedVac and reconstituted in solution A (2% acetonitrile, 1% TFA) and centrifuged (18,000 ×*g*, 10 min).

Peptides were separated on a 15-cm C18 EasySpray column (ThermoFisher ES904, 30°C) after loading onto a 2-cm C18 trap, using a 70-min 10–60% Buffer B (80% acetonitrile, 0.1% formic acid) gradient at 250 nL/min, and analysed on an Orbitrap Exploris (ThermoFisher) with FAIMS Pro™ (CV −45 V) in Data-Independent Acquisition (DIA) mode. Full MS scans (400–1000 m/z, 120,000 resolution, AGC 300%, auto IT) were followed by DIA MS2 scans (60,000 resolution, AGC 1000%, auto IT, NCE 32) using 6-m/z isolation windows with 1-m/z overlap, covering 400–1000 m/z across three consecutive DIA experiments (400–600, 600–800, 800–1000 m/z). Instrument performance was verified using complex lysate QC standards. Proteomic sample analysis was performed at the Technical University of Denmark through Biogenity. EV fractions included in the analysis were isolated from five animals (three males, two females) per age and genotype. No samples were excluded.

#### MS biostatistics data analysis

Raw DIA-MS data were processed with SpectronautTM (v19.1) against the *Mus musculus* UniProt database (taxonomy ID: 10090). Downstream analyses were conducted in R (v4.4.2) using Bioconductor 3.20 packages.[3] To ensure the robustness of the findings, two independent research teams applied two different and blinded methodologies to analyse this dataset.

Regarding one of these methodologies, to handle left-censored missing values in label-free EV proteomics while preserving biologically significant zeros, a bifurcated hybrid statistical pipeline was implemented. For quantitative assessment, proteins with at least four valid intensity values were retained, imputed with the MinProb method ($q = 0.01$) and normalized using variance stabilizing normalization (VSN) to correct mean-variance dependencies. Differential abundance was evaluated with multifactorial linear models in limma package using empirical Bayes moderation to shrink variance estimates. In parallel, proteins completely undetected in one group but quantified in ≥80% of replicates in another were treated as Missing Not At Random (MNAR) in that given pairwise comparison. These proteins were assigned a global minimum limit of detection, and differential presence/absence was robustly assessed using pairwise Fisher's exact tests on binarized detection matrices, with Fold Changes (FC) derived from the VSN-imputed matrix. False Discovery Rate (FDR) was controlled across both analytical branches by local empirical null modelling (fdrtool), and proteins were considered significantly altered at adjusted p-value $< 0.05$ and a FC $> 1.5$. Sample clustering and group similarity were explored through Principal Component Analysis (PCA) with 95% confidence ellipses, and co-expression modules were identified by k-means clustering of row-centred significant proteins. Euclidean distance was used to characterize global proteomic variation. An additive PERMANOVA model incorporating EV fraction, age, and genotype was fitted using marginal tests and 9,999 unrestricted permutations to quantify the multivariate variation independently associated with each factor. To compare genotype-related separation across EV fractions, Euclidean distances between the $APP^{KI}$ and WT multivariate centroids were calculated separately for mitovesicles and sEVs at 2 and 12 months. At each age, the difference between the $APP^{KI}$-WT centroid distances for mitovesicles and sEVs was evaluated using a stratified permutation test. The permutation-derived null distribution was generated by randomly reallocating genotype labels within each EV-fraction stratum while preserving the observed group sizes. Two-sided permutation P values were calculated as the proportion of permuted absolute statistics that were at least as large as the observed absolute statistic, with a plus-one correction. Multiplicity across the two prespecified age-specific distance comparisons was controlled using Holm's procedure.

For the second methodology, a protein co-expression network was built using the protein abundance values, pre-processed as described above (normalized, imputed log2 intensities of the retaine proteins). Pairwise Pearson's correlation coefficients were calculated between all protein pairs across all 20 mitovesicle samples, irrespective of genotype and age. This resulted in a symmetric protein x protein matrix, in which negative correlations and self-correlations were set to zero. Protein modules were identified by maximizing weighted modularity (Q) using the Louvain algorithm,[4] as implemented in the BRAPH toolbox.[5,6] Because Louvain is stochastic, it was run repeatedly (100 runs) and the partition with highest Q among all runs was selected.

To remove weak and noisy connections, correlations below a threshold $\tau$ were set to zero. To select the threshold, the optimal partition ($Q_{max}$) was calculated at each threshold in the range 0.30 to 0.90 (steps of 0.05). At each threshold, a null distribution of Q values was generated by randomly permuting edge weights across all protein pairs (100 permutations), which preserves the number of edges and distribution of edge-weights, but randomizes the correlation structure,[5] and then calculating the optimal partition for each of the randomized null network ($Q_{null}$). The observed $Q_{max}$ was expressed as a z-score relative to the mean and standard deviation of $Q_{null}$, and the threshold with the highest z-score was selected ($\tau = 0.30$).

To obtain a stable partition, Louvain algorithm ($\gamma = 1$) was run 100 times on the weighted network at $\tau = 0.30$. A co-assignment matrix was computed, in which each entry was the fraction of runs in which two proteins were assigned to the same module.[7] Entries below 0.5 were set to zero and self-connections were removed; finally, Louvain algorithm was applied once to this matrix to obtain the final modules.

For each module of interest, protein abundances were z-scored across the 20 mitovesicle samples and PCA was performed on the resulting sample x protein matrix. The first principal component (PC1) scores were used as the eigengene, with the sign set to correlate positively with the mean z-scored abundance of the module's proteins, so higher values indicate higher module abundance. Four comparisons (WT 2 vs 12 months; $App^{KI}$ 2 vs 12 months; WT vs $App^{KI}$ at 2 months; WT vs $App^{KI}$ at 12 months) were tested per module with two-sample t-test; P values were adjusted within each module by controlling the false discovery rate.[8]

Functional characterization was performed using over-representation analysis (ORA) and gene set enrichment analysis (GSEA) with clusterProfiler and fgsea, ranking proteins by log2FC x -log10(p-value). Query databases included KEGG, Gene Ontology (Biological Processes, Molecular Function, Cellular Component), and Reactome, plus custom annotation sets for subcellular localisation (Human Protein Atlas, UniProt, Hein *et al.* 2025) and for submitochondrial localisation and MitoPathways (MitoCarta 3.0).[9-16] Mouse and human protein identifiers (Symbol, UniProt ID, and Entrez ID) were mapped using mapIds, homologene, and manual curation. To reduce redundancy, significant terms were collapsed into representative meta-clusters based on GOSemSim or Jaccard similarity indices and dynamic tree cutting (dynamicTreeCut). Cluster representative keywords were subsequently extracted using natural language processing tokenization.

#### Human proteomic dataset and cross-species analysis

The original mass spectrometry proteomics data has been published[17] and can be found on the ProteomeXchange Consortium via the PRIDE partner repository with the dataset identifier PXD037708. Briefly, the authors analysed postmortem frontotemporal cortex and hippocampus from eight donors (aged 47-89+), all diagnosed with AD at autopsy and classified as A3, B3, C3 (with one case A3, B3, C2) according to the National Institute on Aging-Alzheimer's Association guidelines.[18] EVs were isolated by density-gradient centrifugation into eight fractions, with F7–8 enriched in mitochondrial proteins and consistent with mitovesicles, using the same isolation protocol used for murine vesicles in this study.[1] Ethical approval, tissue collection and informed consent procedures are described in the original publication. No human tissues were collected or analysed in the present study.

To evaluate cross-species conservation of the mitovesicle-specific proteomic signature, the murine EV dataset was integrated with the human AD EV cohort. Consistent with our initial analytical framework, two research teams performed this cross-species evaluation independently using different methodologies.

In the first approach, Human UniProt identifiers were mapped to murine orthologs using homologene, supplemented by manually curated Entrez/UniProt dictionaries for unmapped IDs. The integrated abundance matrix underwent MinProb imputation and joint VSN to simultaneously mitigate batch effects and cross-study technical variance. For the quantitative branch, proteins were retained if detected in at least six human and four mouse samples before imputation, while MNAR evaluations kept the condition-specific 80% detection requirement. Differential analysis followed the aforementioned hybrid pipeline, with stringent thresholds (FDR $< 0.001$, linear FC $> 5$) to isolate high-confidence conserved targets. Conserved cross-species mitovesicle markers were ultimately defined by intersecting murine and human k-means co-expression clusters, and ORA and GSEA were performed as described previously.

In the second methodology, we compared 12-month old $APP^{KI}$ mitovesicles and sEVs (n = 5 for each) with human AD fractions F7 and F5 (n = 8 for each), restricting the analysis to the 2,348 proteins quantified in all 26 samples. Within each group, log2 transformed protein abundancies were averaged and proteins were ranked from lowest (rank 1) to highest (rank 2,348), with ties receiving average ranks. For each protein, we calculated (i) the rank discrepancy between human F7 and mouse mEV, |rank F7 − rank mEV|; (ii) the signed mouse rank difference (rank mEV − rank sEV); and (iii) the signed human rank difference (rank F7 − rank F5). Proteins were assigned to the conserved mitovesicle signature if they met all three criteria: concordant rank across species (lowest quartile of (i)), higher rank in mitovesicles than sEVs (highest quartile of (ii)), and higher rank in F7 than F5 (highest quartile of (iii)). In total, 105 proteins satisfied all criteria. For visualization (Fig. 6), marker size was scaled linearly with the combined score z(mEV − sEV) + z(F7 − F5) − z(|F7 − mEV|), where z denotes the z-score across all proteins.

Subsequently, the results from both approaches were cross compared to identify the overlapping proteomic profile and robustly validate the findings.

### Western blotting

sEVs and mitovesicles isolated from three individual mice or total forebrain homogenates from 12-month-old mice (*n* = five/group), females only, were lysed using RIPA buffer supplemented with protease inhibitors. Protein concentration was determined by Bradford colorimetric assay. Samples containing 10–20 µg of protein were denatured in sample buffer for 5 min at 95ºC and then loaded with the ECL™ Rainbow™ Marker on NuPAGE 4–12% Bis-Tris gels in MES SDS running buffer. Proteins were transferred onto 0.45 µm nitrocellulose membranes for 2 h. The membranes were stained with Ponceau S to enable normalization to total protein content before being blocked for 1 h at 21°C with 5% BSA/TBS-T. Afterwards, membranes were incubated overnight at 4°C with primary antibodies diluted in 5% BSA/TBS-T (**Supplementary table 1**). Before developing, membranes were washed and incubated with the appropriate secondary antibodies for 1 h at 21°C (**Supplementary table 1**). Odyssey CLx Imaging System (LICORBio) was used to develop the membranes, and image quantification was performed in Fiji.

#### Mitovesicle respiratory capacity by Seahorse Analyser

Seahorse is a technique that allows the measurement of oxygen consumption rate (OCR) in live cells or isolated mitochondria, thus providing information on bioenergetics.

To assess the respiratory capacity, mitovesicles were isolated from eight individual mice (three males, five females) per group, run in duplicates or triplicates. Fifteen µg of mitovesicles were then resuspended in 30 µL mitochondria assay solution (MAS; 70 mM sucrose, 220 mM mannitol, 10 mM $KH_2PO_4$, 5 mM $MgCl_2$, 1 mM EGTA, 2 mM HEPES–KOH, pH 7.2) supplemented with 0.2% fatty acid-free BSA and 8% polyethylene glycol to precipitate mitovesicles to the bottom of the plate for optimal readout. Mitovesicles were transferred to a poly(ethyleneimine)-coated (1:15,000) Agilent 96-well Seahorse XF96 microplate and centrifuged at 2,200 ×*g* for 20 min at 4°C to pellet the mitovesicles at the bottom of the plate. The day before the experiment, the Seahorse XF96 sensor cartridge (Agilent) was hydrated and calibrated according to the manufacturer's instructions. OCR was measured using the Seahorse XF96 analyser (Agilent).

To evaluate the individual mitochondrial complexes activity, MAS was further supplemented with 0.1 mM ADP-K (Complex V substrate), 10 mM pyruvate (Complex I substrate), 2 mM malate (Complex I substrate), pH 7.4, and 4 µM FCCP (Carbonyl cyanide 4-(trifluoromethoxy)phenylhydrazone; mitochondrial uncoupler). Pyruvate and malate provide NADH-linked substrates for Complex I-supported respiration, whereas FCCP dissipates the proton gradient, allowing measurement of maximal electron transport capacity. OCR was measured at baseline and after sequential stimulation with 2 µM rotenone (Complex I inhibitor), 10 mM succinate (Complex II substrate), 4 µM antimycin A (Complex III inhibitor) and 10 mM ascorbate/0.1 mM TMPD (N,N,N′,N′-tetramethyl p-phenylenediamine dihydrochloride; electron donors to cytochrome C/complex IV) as previously described.[19] Accordingly, NADH-linked basal respiration and individual complex-dependent respiration were calculated based on the OCR. Data are given in pmol $O_2$/min.

#### Statistical analysis

Statistical analysis was done using GraphPad Prism version 10 and data are presented as mean ± SEM. n refers to the number of unique individual biological samples. No blinding nor randomization of samples were performed. Data distribution was assessed for normality, and appropriate parametric or non-parametric tests were applied, including unpaired t-tests and one-way ANOVA or 2way ANOVA followed Šídák's multiple comparisons test. Outliers were identified using the ROUT method Q = 2%). A $P < 0.05$ threshold was considered statistically significant. $P$ values shown for NTA are adjusted $P$ values.

### Supplementary figures and tables

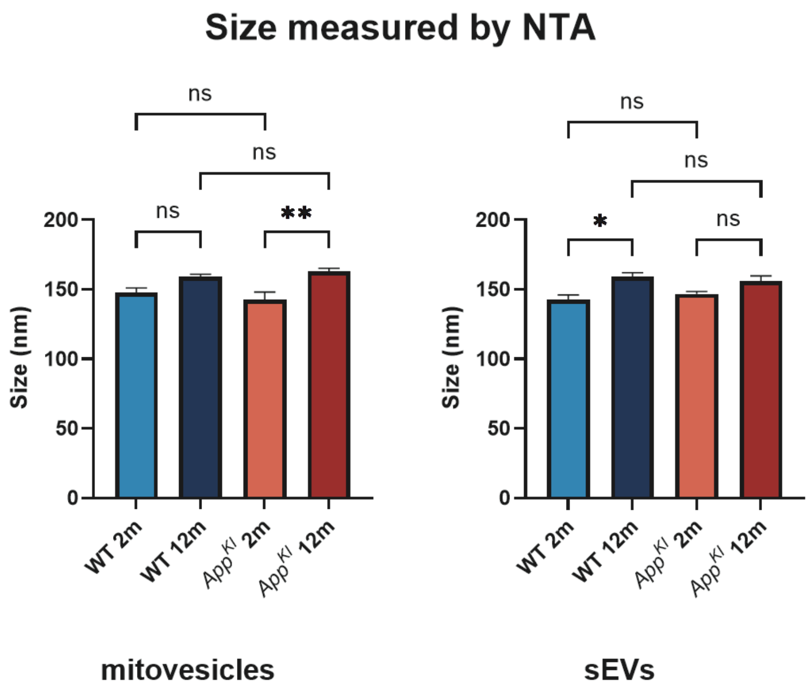


**Supplementary Figure 1. Vesicle size measured by NTA.** Quantification of size of each vesicle population measured by NTA. Data presented as mean ± SEM and statistical significance was done using one-way ANOVA. * $P \leq 0.05$, ** $P \leq 0.01$, ns. non-significant.

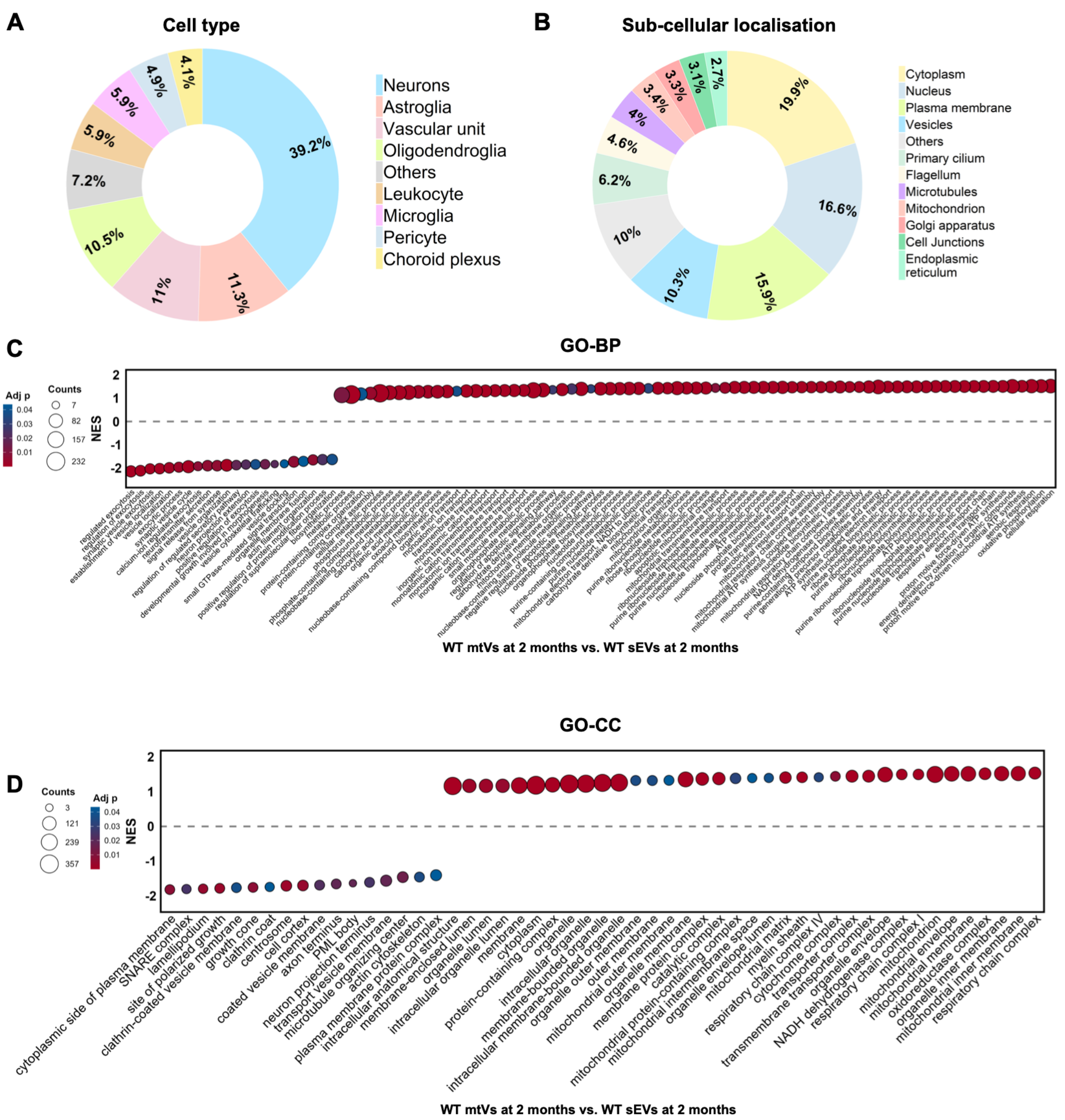


**Supplementary Figure 2. Proteomic characteristics of sEVs and comparison with mitovesicles in 2-month-old WT brain.** (**A**–**B**) Pie charts depicting the inferred cell-type origin (**A**) and subcellular organization (**B**) of proteins detected in WT sEV fraction at 2 months. (**C**–**D**) Dot plots showing the full GSEA of GO-BP terms (**C**) and GO-CC terms (**D**) of mitovesicles vs. sEVs in 2-month-old WT brain.

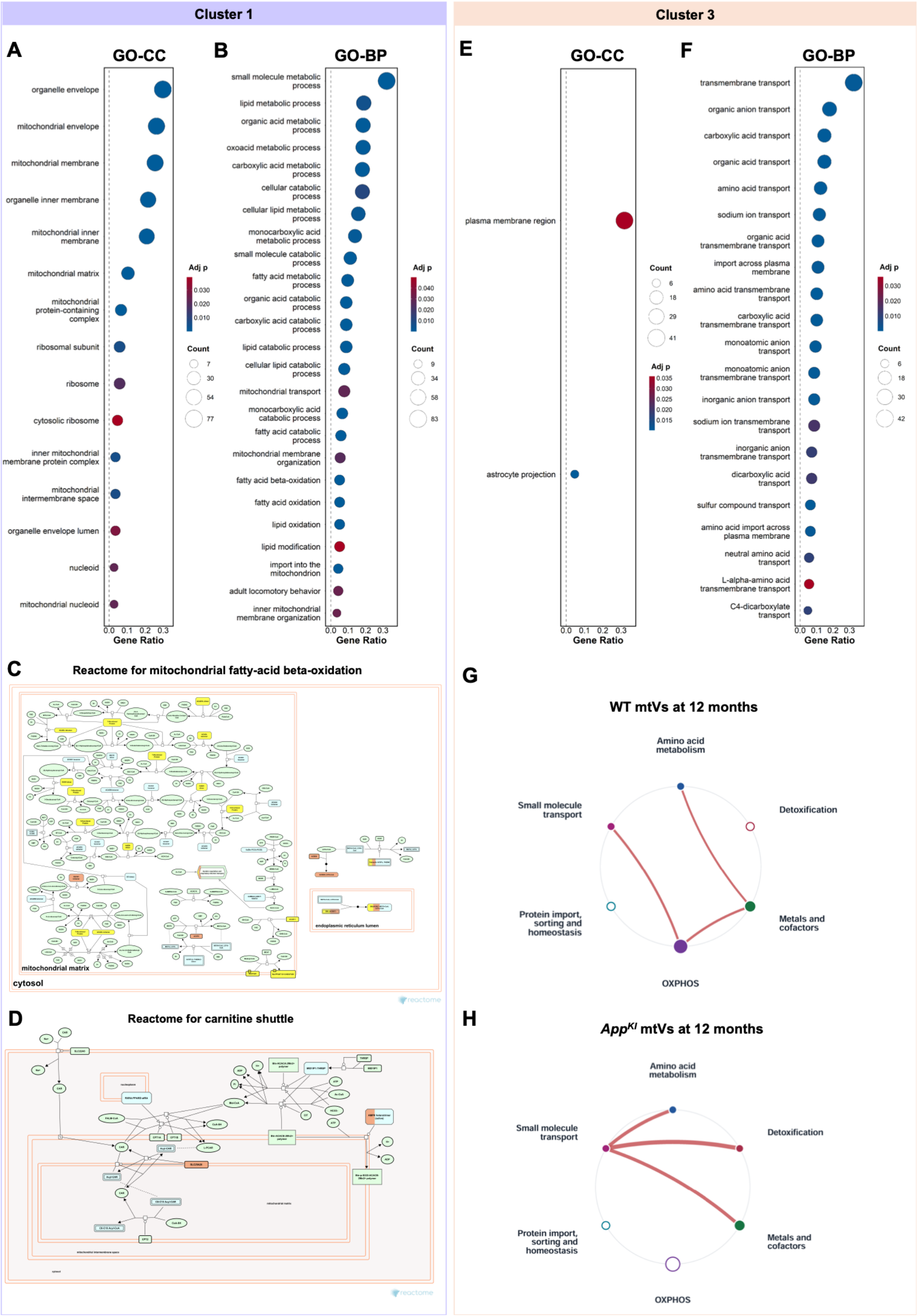


**Supplementary Figure 3. Deeper analyses of Clusters 1 and 3.** (**A**–**B**) Dot plots showing ORA of Cluster 1 enriched GO-CC (**A**) and GO-BP (**B**) terms. (**C**–**D**) Reactome pathways of the mitochondrial β-oxidation (**C**) and carnitine shuttle (**D**). Yellow labelled proteins are present in Cluster 1, orange labelled proteins were found in mitovesicles but not in Cluster 1. (**E**–**F**) Dot plots showing the ORA of Cluster 3 enriched GO-CC (**E**) and GO-BP (**F**) terms. (**G**–**H**) Network analysis of the relationships between mitochondrial processes in mitovesicles from 12-month-old WT **(G)** and App$^{KI}$ 12-month-old **(H)** mice. Nodes represent MitoPathways and edges represent the average Pearson correlation between the proteins of two pathways calculated across samples (*n* = 5 per group). Edge thickness is proportional to correlation strength and node size reflects the number of proteins within the corresponding process. For visualization, correlations below 0.8 were set to zero. Nodes without remaining connections are shown as open circles.

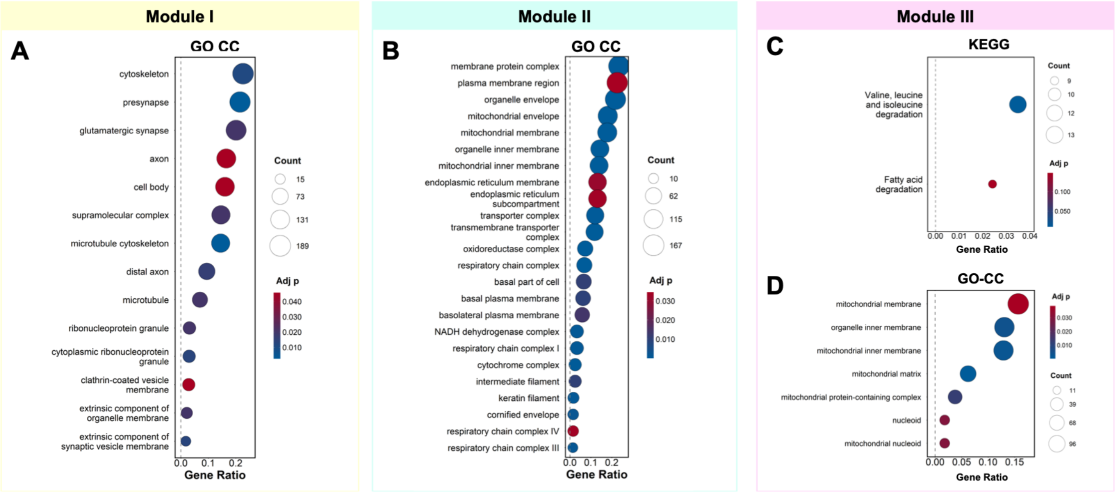


**Supplementary Figure 4**. **ORA of Modules I-III**. (**A**, **B**) Dot plots showing ORA of GO-CC terms for Module I (**A**) and II (**B**). (**C**, **D**) Dot plots showing ORA of KEGG pathways (**C**) and GO-CC terms (**D**) of Module III.

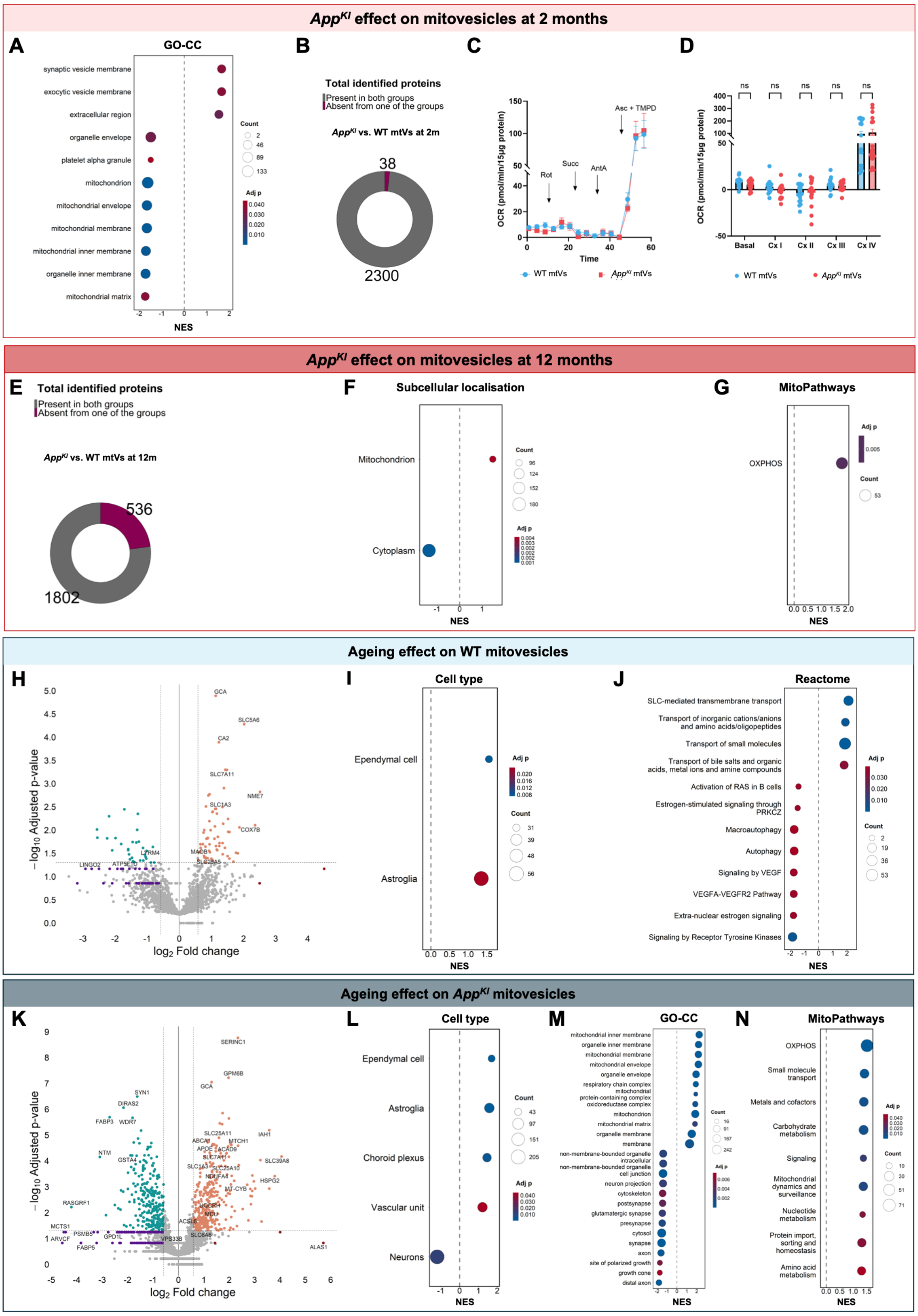


**Supplementary Figure 5**. **Disease- and age-related proteomic shift and functional changes in mitovesicles.** (**A**–**D**) Analyses done in 2-month-old WT vs. *App*$^{KI}$ mitovesicles. (**A**) Dot plot representing GSEA of GO-CC terms. (**B**) Donut plot showing present and missing proteins in 2-month-old mitovesicles. (**C**) Seahorse uncoupled OCR traces of 2-month-old mitovesicles following sequential injections of rotenone (2 µM), succinate (10 mM), antimycin A (4 µM) and ascorbate (1 mM)/TMPD (100 mM) ($n$ = 17–19). Timing of the stimulators/inhibitors indicated on the graph. (**D**) Bar graph showing respiration of each individual complex ($n$ = 17–19). Data presented as mean ±

SEM and statistical significance was done using one-way ANOVA. ns. non-significant. (**E**–**G**) Analyses done in 12-month-old WT vs. *App*$^{KI}$ mitovesicles. (**E**) Donut plot showing present and missing proteins. (**F**–**G**) Dot plots representing subcellular organization (**F**) and MitoPathways (**G**). (**H**–**J**) Pairwise analyses of presymptomatic vs. symptomatic mitovesicles isolated from WT mice. (H) Volcano plot. (**I, J**) Dot plots showing cell-type distribution (**I**) and Reactome pathway analysis (**J**). (**K**–**N**) Pairwise analyses of presymptomatic vs. symptomatic mitovesicles isolated from *App*$^{KI}$ mice. (**K**) Volcano plot. (**L**–**N**) Dot plots showing cell-type distribution (**L**), ORA of GO-CC terms (**M**) and MitoPathways (**N**).

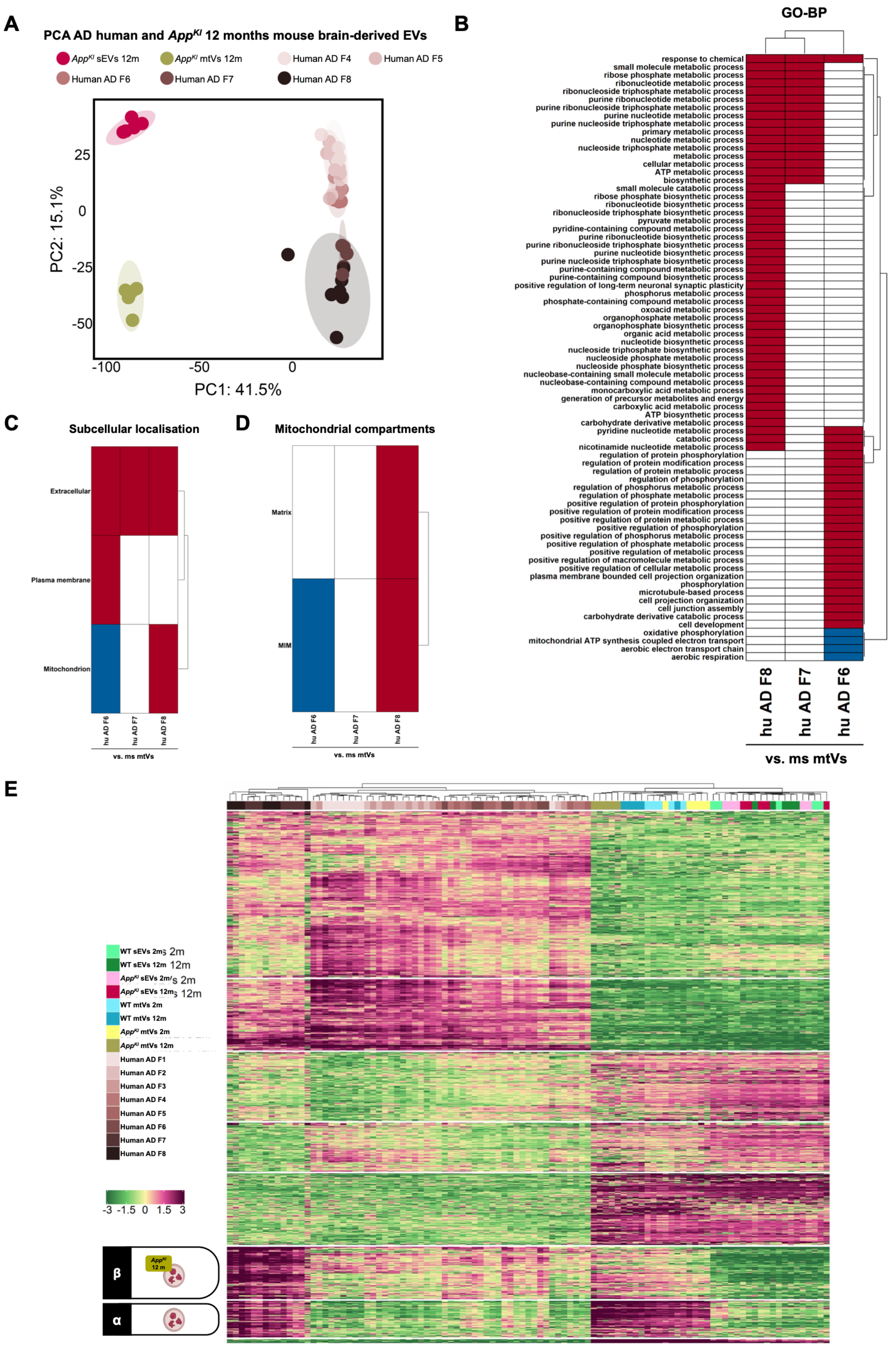


**Supplementary Figure 6. Murine AD mitovesicles in comparison with human AD patient mitovesicles.** (**A**) PCA of human sEVs (AD F4–6), human mitovesicles (AD F7–8) and symptomatic mouse sEVs and mitovesicles. (**B–D**) Comparison of pathway representation in human fractions F6-8 relative to the mouse mitovesicle proteome for GO-BP (**B**), subcellular localization (**C**) and mitochondrial compartments (**D**). Red indicates higher representation; blue indicates lower representation and white indicates comparable representation. (**E**) Heatmap showing hierarchical clustering of murine and human EV proteomes. Cluster α indicated mitovesicle proteins in human AD samples. Cluster β mitovesicle proteins affected by amyloid pathology.

**Table 1. List of antibodies for Western blot**

| Antibody | Dilution | Brand | Catalogue number |
|---|---|---|---|
| **Primary antibodies** | | | |
| GAP43 | 1:500 | Proteintech | 16971-1-AP |
| OXPHOS | 1:1000 | Abcam | ab110413 |
| NDUFB3 | 1:200 | Abcam | ab202585 |
| VDAC1/2/3 | 1:1000 | Abcam | ab15895 |
| ACAD9 | 1:1000 | Cell Signaling Technology | 9796 |
| FABP3 | 1:1000 | Cell Signaling Technology | 92185 |
| APOE | 1:500 | Invitrogen | 701241 |
| CD81 | 1:500 | Cell Signaling Technology | 10037 |
| Flotillin-1 | 1:500 | BD Biosciences | 610820 |
| **Secondary antibodies** | | | |
| IRDye 800CW anti-mouse | 1:20,000 | LICORBio | 926-32212 |
| IRDye 800CW anti-rabbit | 1:20,000 | LICORBio | 926-32213 |
| IRDye 680RD anti-mouse | 1:20,000 | LICORBio | 926-68072 |
| IRDye 680RD anti-rabbit | 1:20,000 | LICORBio | 926-68071 |

**Table 2. List of reagents and materials**

| Reagent | Brand | Catalogue number |
|---|---|---|
| Hibernate-A medium | ThermoFisher | A1247501 |
| Papain | Worthington | LK003176 |
| PMSF | Sigma-Aldrich | P7626 |
| Cell strainer 40 µm | VWR | 76327-098 |
| Syringe filter 0.22 µm | Cytiva | 10462200 |
| OptiPrep™ | StemCell Technologies | 07820 |
| NuPAGE 12% Bis-Tris | Invitrogen | NP0341BOX |
| EZBlue™ Gel Staining Reagent | Sigma | G1041 |
| Pierce BCA Protein Assay Kit | ThermoFisher | 23225 |
| NuPAGE 4–12% Bis-Tris | Invitrogen | NP0321BOX |
| NuPAGE MES SDS running buffer | Invitrogen | NP0002 |
| ECL™ Rainbow™ Marker – Full Range | GE HealthCare | RPN800E |
| Nitrocellulose membranes 0.45 µm | GE Healthcare | 10600002 |
| Ponceau S | Sigma-Aldrich | P7170 |
| Bovine Serum Albumin | Sigma-Aldrich | A6003 |
| Polyethylene glycol (PEG600) | Sigma-Aldrich | 81260 |
| poly(ethyleneimine) | Sigma-Aldrich | 03880 |
| ADP-K | Sigma-Aldrich | A2754 |
| pyruvate | Sigma-Aldrich | 58636 |
| Malate | Sigma-Aldrich | 46940-U |
| FCCP | Sigma-Aldrich | C2920 |
| Rotenone | Sigma-Aldrich | R8875 |
| Succinate | Sigma-Aldrich | S3674 |
| Antimycin A | Sigma Aldrich | A8674 |
| Ascorbate | Sigma-Aldrich | A4403 |
| TMPD | Sigma-Aldrich | T3134 |

**Table 3. Proteins in Cluster 4, 5 and 6**

| Protein | Cluster |
|---|---|
| AIDA | 4 |
| ABCA13 | 4 |
| CUL4B | 4 |
| HOOK3 | 4 |
| APRT | 4 |
| SRGAP2 | 4 |
| ARHGAP44 | 4 |
| SBDS | 4 |
| APLP1 | 4 |
| CTNND1 | 4 |
| RARS1 | 4 |
| GPS1 | 4 |
| STX5 | 4 |
| RRAS | 4 |
| PLS3 | 4 |
| CSE1L | 4 |
| TTYH2 | 4 |
| ADD2 | 4 |
| CADM4 | 4 |
| PPP1CC | 4 |
| PSMA1 | 4 |
| CCDC158 | 4 |
| NRGN | 4 |
| HRAS | 4 |
| SCAMP5 | 4 |
| AHSA1 | 4 |
| ACYP1 | 4 |
| VBP1 | 4 |
| SEPTIN4 | 4 |
| PHYHIP | 4 |
| ARL3 | 4 |
| PPME1 | 4 |
| GRHPR | 4 |
| PCLO | 4 |
| LRPAP1 | 4 |
| SMPD1 | 4 |
| ARMC8 | 4 |
| ELMO1 | 4 |
| LINGO2 | 4 |
| LINGO1 | 4 |
| CYRIB | 4 |
| CADPS2 | 4 |
| KCTD8 | 4 |
| MAP6 | 4 |
| VAT1 | 4 |

| | |
|---|---|
| ATP6V1E1 | 4 |
| PGM2L1 | 4 |
| VAT1L | 4 |
| HPRT1 | 4 |
| MAP1LC3A | 4 |
| LRRC47 | 4 |
| MARCKS | 4 |
| FMN2 | 4 |
| GRM1 | 4 |
| SYT17 | 4 |
| CLTA | 4 |
| CALB2 | 4 |
| GNG7 | 4 |
| GNG2 | 4 |
| MACF1 | 4 |
| GAP43 | 4 |
| BASP1 | 4 |
| APOB | 4 |
| ASAP1 | 4 |
| MBP | 4 |
| GET1 | 4 |
| CSNK1G1 | 4 |
| ERP29 | 4 |
| CRIP1 | 4 |
| CACNB1 | 4 |
| RETREG2 | 4 |
| ATP6V1D | 4 |
| HS2ST1 | 4 |
| KCTD12 | 4 |
| NARS1 | 4 |
| PRKCB | 4 |
| NCALD | 4 |
| CYFIP2 | 4 |
| PLXNA1 | 4 |
| RAP1GDS1 | 4 |
| HPCA | 4 |
| VSNL1 | 4 |
| HPCAL4 | 4 |
| HSPA8 | 4 |
| CHP1 | 4 |
| HPCAL1 | 4 |
| GPI | 4 |
| CLTC | 4 |
| DPYSL2 | 4 |
| AP2A2 | 4 |
| ANXA6 | 4 |
| ATP6V1H | 4 |

| | |
|---|---|
| PLXNC1 | 4 |
| GET3 | 4 |
| AMPD2 | 4 |
| GNAI1 | 4 |
| LDHA | 4 |
| GAS7 | 4 |
| EPB41L3 | 4 |
| RAB3A | 4 |
| AP2A1 | 4 |
| OTUB1 | 4 |
| SNAP91 | 4 |
| DNM1 | 4 |
| OXSR1 | 4 |
| CAMKV | 4 |
| NSF | 4 |
| PRKAR2B | 4 |
| PDE2A | 4 |
| PDE1B | 4 |
| CAB39 | 4 |
| SLC18A2 | 4 |
| CACNA2D1 | 4 |
| WASF1 | 4 |
| SCN2B | 4 |
| PCBD1 | 4 |
| RHOF | 4 |
| ROCK2 | 4 |
| CLTB | 4 |
| SERPINI1 | 4 |
| SNAP29 | 4 |
| CXADR | 4 |
| CUTA | 4 |
| SUSD6 | 4 |
| PLCL2 | 4 |
| UBE2L3 | 4 |
| CCT2 | 4 |
| PDPK1 | 4 |
| CLASP2 | 4 |
| YARS1 | 4 |
| USP14 | 4 |
| MPP2 | 4 |
| MADD | 4 |
| LRRC57 | 4 |
| DMXL2 | 4 |
| UBE2N | 4 |
| HSPA12A | 4 |
| FLOT1 | 4 |
| CCT7 | 4 |

| | |
|---|---|
| KIF2A | 4 |
| GAD1 | 4 |
| CDK5 | 4 |
| CLIP2 | 4 |
| IQCB1 | 4 |
| CLUH | 4 |
| CUL2 | 4 |
| TSC2 | 4 |
| PIP4K2A | 4 |
| CAMK1 | 4 |
| SCRN3 | 4 |
| STK11 | 4 |
| HAGH | 4 |
| MAPRE3 | 4 |
| MYO18A | 4 |
| SH3GLB2 | 4 |
| SRCIN1 | 4 |
| PSMB7 | 4 |
| RHEB | 4 |
| RAB27B | 4 |
| TPPP | 4 |
| TBC1D24 | 4 |
| CSNK1A1 | 4 |
| ACACA | 4 |
| AMPD3 | 4 |
| DPY19L1 | 4 |
| HDHD2 | 4 |
| DIAPH1 | 4 |
| VANGL2 | 4 |
| DCTN2 | 4 |
| BAG6 | 4 |
| INPP4A | 4 |
| MTOR | 4 |
| LIN7C | 4 |
| MARK2 | 4 |
| ATRN | 4 |
| PCSK1N | 4 |
| PRKAG2 | 4 |
| CORO2B | 4 |
| PDCD6IP | 4 |
| TALDO1 | 4 |
| STIP1 | 4 |
| RAB11B | 4 |
| DCTN3 | 4 |
| CORO1C | 4 |
| PPP2R5C | 4 |
| ACTR10 | 4 |

| | |
|---|---|
| SPARCL1 | 4 |
| IDH1 | 4 |
| MAGI1 | 4 |
| RASGRF2 | 4 |
| FBXO22 | 4 |
| TUBA4A | 4 |
| PODXL2 | 4 |
| PTPA | 4 |
| TKFC | 4 |
| NSDHL | 4 |
| KIF5C | 4 |
| DCTN1 | 4 |
| IPO4 | 4 |
| DDT | 4 |
| ORC5 | 4 |
| PSMD8 | 4 |
| RAPGEF2 | 4 |
| UBE3C | 4 |
| KCNIP3 | 4 |
| PPM1E | 4 |
| CBR1 | 4 |
| PPP3CA | 4 |
| IPO5 | 4 |
| DNM3 | 4 |
| SNX4 | 4 |
| ACAP2 | 4 |
| PDCD5 | 4 |
| FCSK | 4 |
| KLC1 | 4 |
| UBE3B | 4 |
| PFDN2 | 4 |
| PHGDH | 4 |
| UGP2 | 4 |
| PSMC4 | 4 |
| FBXL16 | 4 |
| USP5 | 4 |
| NPEPPS | 4 |
| PPP2R1A | 4 |
| ENO1 | 4 |
| MARK1 | 4 |
| TOM1 | 4 |
| NAP1L4 | 4 |
| EPHB3 | 4 |
| ZZEF1 | 4 |
| PPP2R5A | 4 |
| MATK | 4 |
| ELP3 | 4 |

| | |
|---|---|
| PEF1 | 4 |
| ENPP4 | 4 |
| SNF8 | 4 |
| CYRIA | 4 |
| STK39 | 4 |
| PSMB3 | 4 |
| IPO9 | 4 |
| TRIO | 4 |
| AP1S1 | 4 |
| PFKL | 4 |
| CKAP5 | 4 |
| XPO1 | 4 |
| CDC37 | 4 |
| KIF21A | 4 |
| EPHA4 | 4 |
| PRKAG1 | 4 |
| GIT1 | 4 |
| SCAI | 4 |
| MTMR9 | 4 |
| ACTR3B | 4 |
| TUBGCP3 | 4 |
| MAPK3 | 4 |
| SKT | 4 |
| XPO7 | 4 |
| RASAL1 | 4 |
| PPP5C | 4 |
| DAAM1 | 4 |
| PSMA3 | 4 |
| CAPZB | 4 |
| TTC7B | 4 |
| GSTA4 | 4 |
| TWF2 | 4 |
| EFR3B | 4 |
| HSPA4L | 4 |
| PSD3 | 4 |
| AP3D1 | 4 |
| IPO7 | 4 |
| WDR7 | 4 |
| KPNB1 | 4 |
| NDRG3 | 4 |
| PSMC1 | 4 |
| PICALM | 4 |
| UBQLN1 | 4 |
| PLCXD3 | 4 |
| PSMA7 | 4 |
| STXBP5 | 4 |
| KCNAB2 | 4 |

| | |
|---|---|
| RAP2A | 4 |
| RAP2B | 4 |
| RAB5C | 4 |
| GNG13 | 4 |
| ADCY1 | 4 |
| PIP5K1C | 4 |
| SEPTIN8 | 4 |
| NUCB1 | 4 |
| ROGDI | 4 |
| PLXND1 | 4 |
| RUFY3 | 4 |
| HTT | 4 |
| RWDD1 | 4 |
| SBF1 | 4 |
| NMT1 | 4 |
| CALB1 | 4 |
| OSBPL1A | 4 |
| RGS7BP | 4 |
| PDXK | 4 |
| NDRG4 | 4 |
| ABRACL | 4 |
| BMERB1 | 4 |
| PEPD | 4 |
| STUB1 | 4 |
| ICAM5 | 4 |
| FLOT2 | 4 |
| FDPS | 4 |
| ATP6V1A | 4 |
| ATP6V1G2 | 4 |
| RGS7 | 4 |
| SEPTIN3 | 4 |
| STXBP1 | 4 |
| SEPTIN7 | 4 |
| CADPS | 4 |
| UFC1 | 4 |
| AP1B1 | 4 |
| PACSIN1 | 4 |
| NAPB | 4 |
| EXTL2 | 4 |
| HSPA4 | 4 |
| NCS1 | 4 |
| ATP6V1C1 | 4 |
| KRAS | 4 |
| CAND1 | 4 |
| CUL3 | 4 |
| GSTO1 | 4 |
| RAB5A | 4 |

| | |
|---|---|
| UBA1 | 4 |
| NAA15 | 4 |
| PSMD13 | 4 |
| BRK1 | 4 |
| ADD1 | 4 |
| RFTN2 | 4 |
| YWHAQ | 4 |
| DNM1L | 4 |
| NBEA | 4 |
| CSNK2A2 | 4 |
| PI4KA | 4 |
| EEF1A2 | 4 |
| CCT4 | 4 |
| SYT3 | 4 |
| LANCL2 | 4 |
| GABBR2 | 4 |
| PACSIN2 | 4 |
| TMX2 | 4 |
| VPS45 | 4 |
| ATP2B1 | 4 |
| GUCY1B1 | 4 |
| GRM5 | 4 |
| IDE | 4 |
| ANO6 | 4 |
| GRID1 | 4 |
| CARS1 | 4 |
| MPI | 4 |
| JAKMIP3 | 4 |
| PTPRN2 | 4 |
| WDR47 | 4 |
| KCNA1 | 4 |
| ITGAM | 4 |
| KCNA4 | 4 |
| EHD1 | 4 |
| CPLX1 | 4 |
| PLXNA2 | 4 |
| GSK3A | 4 |
| PRUNE2 | 4 |
| PI4K2A | 4 |
| PIK3R4 | 4 |
| CADM3 | 4 |
| ARFGEF1 | 4 |
| MAP2K1 | 4 |
| ARHGEF5 | 4 |
| LMF1 | 4 |
| GNB1 | 4 |
| ACLY | 4 |

| | |
|---|---|
| COPS2 | 4 |
| EXOC4 | 4 |
| KIF1A | 4 |
| PPFIA3 | 4 |
| INPP1 | 4 |
| SGSM1 | 4 |
| RAB3GAP2 | 4 |
| SULT4A1 | 4 |
| PTK2 | 4 |
| DYNC1I1 | 4 |
| LIN7A | 4 |
| WNK2 | 4 |
| MPP1 | 4 |
| EPHA5 | 4 |
| AGTPBP1 | 4 |
| PPP6R1 | 4 |
| NECAB1 | 4 |
| NUDCD3 | 4 |
| VPS28 | 4 |
| PDAP1 | 4 |
| ACY1 | 4 |
| ITFG1 | 4 |
| CYTH2 | 4 |
| RAB4A | 4 |
| UCHL5 | 4 |
| EXOC7 | 4 |
| WDR37 | 4 |
| OGFRL1 | 4 |
| MAP2K4 | 4 |
| PSMD11 | 4 |
| PDCD10 | 4 |
| TLN2 | 4 |
| LTA4H | 4 |
| EXOC5 | 4 |
| SPTB | 4 |
| SACM1L | 4 |
| DOCK9 | 4 |
| NECAP1 | 4 |
| FUT8 | 4 |
| RAB3B | 4 |
| SMAP2 | 4 |
| PTPRE | 4 |
| VPS53 | 4 |
| AP3B2 | 4 |
| GGPS1 | 4 |
| NECAB2 | 4 |
| CAB39L | 4 |

| | |
|---|---|
| CCDC43 | 4 |
| DPP3 | 4 |
| BLTP1 | 4 |
| EIF4E | 4 |
| PPP1CB | 4 |
| CAPN5 | 4 |
| PPP1R9B | 4 |
| KIF5B | 4 |
| TMEM30A | 4 |
| ST8SIA5 | 4 |
| STMN1 | 4 |
| NEDD8 | 4 |
| GNPDA1 | 4 |
| C2CD5 | 4 |
| BLVRA | 4 |
| EXOC8 | 4 |
| BIN1 | 4 |
| SH3GL2 | 4 |
| TMEM68 | 4 |
| GGT7 | 4 |
| TM9SF1 | 4 |
| RHOA | 4 |
| HSPH1 | 4 |
| KCTD16 | 4 |
| AMPH | 4 |
| SARS1 | 4 |
| NME2 | 4 |
| LANCL1 | 4 |
| RGS6 | 4 |
| ARF6 | 4 |
| ST13 | 4 |
| SEPTIN9 | 4 |
| EXOC2 | 4 |
| ATL1 | 4 |
| EHD3 | 4 |
| EPN2 | 4 |
| BZW1 | 4 |
| COPS6 | 4 |
| ARMT1 | 4 |
| UBL4A | 4 |
| CACYBP | 4 |
| EPM2AIP1 | 4 |
| MAP2K6 | 4 |
| IMPA1 | 4 |
| RAB5B | 4 |
| GPD1L | 4 |
| NAPG | 4 |

| | |
|---|---|
| ABI2 | 4 |
| PLXNA4 | 4 |
| PDXP | 4 |
| WARS1 | 4 |
| TSG101 | 4 |
| ABI1 | 4 |
| DYNLRB1 | 4 |
| PDE1A | 4 |
| SCRN1 | 4 |
| ENO2 | 4 |
| PHF24 | 4 |
| ATP6V1G1 | 4 |
| MAN2A2 | 4 |
| ENOPH1 | 4 |
| CAPN1 | 4 |
| PTPN11 | 4 |
| SLITRK1 | 4 |
| AGBL4 | 4 |
| EFR3A | 4 |
| STX6 | 4 |
| GMFB | 4 |
| IQGAP1 | 4 |
| UBL3 | 4 |
| RPTOR | 4 |
| SEC14L2 | 4 |
| RDX | 4 |
| SGTA | 4 |
| RPAP3 | 4 |
| PACS1 | 4 |
| SGPL1 | 4 |
| PAK3 | 4 |
| ITGB2 | 4 |
| ADRA1A | 4 |
| EPHA7 | 4 |
| FNBP1L | 4 |
| AFDN | 4 |
| ADPRH | 4 |
| GOSR1 | 4 |
| DNAJC8 | 4 |
| MME | 4 |
| TPRG1L | 4 |
| HSBP1 | 4 |
| PAK1 | 4 |
| SEC31A | 4 |
| SCAP | 4 |
| SH3GL3 | 4 |
| ARIH1 | 4 |

| | |
|---|---|
| ARHGAP21 | 4 |
| PRKAA1 | 4 |
| COMMD10 | 4 |
| MEGF11 | 4 |
| TARS1 | 4 |
| GSK3B | 4 |
| IMPACT | 4 |
| DYNLL1 | 4 |
| MAP2K7 | 4 |
| GGCT | 4 |
| TMEM151B | 4 |
| VPS50 | 4 |
| SEC61B | 4 |
| ARFGEF2 | 4 |
| OXR1 | 4 |
| TESC | 4 |
| MAPRE2 | 4 |
| COPS5 | 4 |
| ZNF770 | 4 |
| PNPO | 4 |
| UBA3 | 4 |
| SRR | 4 |
| CAPNS1 | 4 |
| MAPT | 4 |
| PSMC5 | 4 |
| MAP2 | 4 |
| TNPO1 | 4 |
| TUBA8 | 4 |
| CNDP2 | 4 |
| PRKAA2 | 4 |
| ZNF365 | 4 |
| ARHGEF12 | 4 |
| OSCP1 | 4 |
| GOLGA7 | 4 |
| PSME1 | 4 |
| USP9X | 4 |
| MAP1LC3B | 4 |
| HGS | 4 |
| MIF | 4 |
| CBL | 4 |
| PSMB6 | 4 |
| DLGAP4 | 4 |
| SYNE1 | 4 |
| GAK | 4 |
| FMNL1 | 4 |
| CCDC177 | 4 |
| TPP2 | 4 |

| | |
|---|---|
| PEX5L | 4 |
| DGKB | 4 |
| ISOC1 | 4 |
| POGLUT1 | 4 |
| TRIQK | 4 |
| ASNS | 4 |
| MINPP1 | 4 |
| ALAD | 4 |
| IPO11 | 4 |
| SLC24A4 | 4 |
| PAK5 | 4 |
| SGIP1 | 4 |
| GPR26 | 4 |
| PGM3 | 4 |
| FIBP | 4 |
| NOS1 | 4 |
| ASL | 4 |
| UPF2 | 4 |
| GBA2 | 4 |
| ZC2HC1A | 4 |
| ADCY9 | 4 |
| AKR1B1 | 4 |
| DCUN1D1 | 4 |
| EHD4 | 4 |
| REEP5 | 4 |
| CFAP65 | 4 |
| FAM234B | 4 |
| PTPRC | 4 |
| PFDN1 | 4 |
| ZWINT | 4 |
| PDXDC1 | 4 |
| RAP2C | 4 |
| LY6H | 4 |
| BCAT2 | 4 |
| EPHB1 | 4 |
| STMN2 | 4 |
| EPHB2 | 4 |
| SNX5 | 4 |
| GPR158 | 4 |
| CYP46A1 | 4 |
| EPHX2 | 4 |
| EPN1 | 4 |
| NTRK2 | 4 |
| PLCH1 | 4 |
| EIF5B | 4 |
| MIER1 | 4 |
| BSG | 4 |

| | |
|---|---|
| MYO15A | 4 |
| RTP3 | 4 |
| CHRM1 | 4 |
| TSN | 4 |
| TPT1 | 4 |
| PATL1 | 4 |
| SLC8A2 | 4 |
| SERPINB6 | 4 |
| WASL | 4 |
| PTS | 4 |
| CRYM | 4 |
| ATIC | 4 |
| EPHA6 | 4 |
| CPPED1 | 4 |
| UBE2O | 4 |
| SCN1A | 4 |
| FBXL20 | 4 |
| GARS1 | 4 |
| EHBP1 | 4 |
| AGFG1 | 4 |
| CEP170 | 4 |
| L1CAM | 4 |
| FBXO3 | 4 |
| TMSB10 | 4 |
| SMS | 4 |
| USP7 | 4 |
| HECTD3 | 4 |
| KSR1 | 4 |
| SERPINA1B | 4 |
| STXBP6 | 4 |
| WNT10B | 4 |
| MEST | 4 |
| PRKAR1B | 4 |
| MCTS1 | 4 |
| SNX24 | 4 |
| OSBPL9 | 4 |
| FABP5 | 4 |
| HNRNPA3 | 5 |
| RPS15 | 5 |
| HNRNPA2B1 | 5 |
| RPS3A | 5 |
| HNRNPA1 | 5 |
| RPL8 | 5 |
| EIF5A | 5 |
| GLRX3 | 5 |
| NCL | 5 |
| RPL18A | 5 |

| | |
|---|---|
| HNRNPU | 5 |
| G3BP1 | 5 |
| PHB1 | 5 |
| F8 | 5 |
| SYNCRIP | 5 |
| ZYG11B | 5 |
| WASHC4 | 5 |
| CHID1 | 5 |
| TM9SF2 | 5 |
| LYSMD2 | 5 |
| VPS13C | 5 |
| SSR1 | 5 |
| NEDD4 | 5 |
| CLIC4 | 5 |
| CXCL11 | 5 |
| GABRA3 | 5 |
| PIP4K2B | 5 |
| CPNE3 | 5 |
| TAGLN3 | 5 |
| LMNA | 5 |
| TXN | 5 |
| CACNA1C | 5 |
| CACNG3 | 5 |
| RPS3 | 5 |
| PA2G4 | 5 |
| GPR61 | 5 |
| GNA12 | 5 |
| SIRPA | 5 |
| SEC61A2 | 5 |
| UCHL1 | 5 |
| CDH9 | 5 |
| QPCTL | 5 |
| GAD2 | 5 |
| FLNA | 5 |
| RPS5 | 5 |
| PPP4R4 | 5 |
| PIK3R1 | 5 |
| TM9SF4 | 5 |
| MANF | 5 |
| CORO1A | 5 |
| GABARAPL2 | 5 |
| SEZ6L2 | 5 |
| UBE2M | 5 |
| DLG5 | 5 |
| SMPD3 | 5 |
| PPP2R2A | 5 |
| PTGES3 | 5 |

| | |
|---|---|
| FYN | 5 |
| HYCC2 | 5 |
| ACTR3 | 5 |
| PDIA6 | 5 |
| ATP1B4 | 5 |
| UGT8 | 5 |
| CCT6A | 5 |
| ERC2 | 5 |
| THOP1 | 5 |
| TMX4 | 5 |
| ELMO2 | 5 |
| NME3 | 5 |
| IGLON5 | 5 |
| ACTR2 | 5 |
| FABP3 | 5 |
| DPYSL3 | 5 |
| CPLX2 | 5 |
| ALDOA | 5 |
| SEPTIN5 | 5 |
| ATP1A1 | 5 |
| CRMP1 | 5 |
| SYN2 | 5 |
| SYN1 | 5 |
| SNX12 | 5 |
| PGK1 | 5 |
| DIRAS2 | 5 |
| DCLK1 | 5 |
| PGRMC1 | 5 |
| RPH3A | 5 |
| RRAS2 | 5 |
| RAB23 | 5 |
| GNB5 | 5 |
| SYT7 | 5 |
| MAP1A | 5 |
| ACOT7 | 5 |
| CMPK1 | 5 |
| PPA1 | 5 |
| PRKAR2A | 5 |
| RAB3C | 5 |
| RAB33B | 5 |
| CYB5D2 | 5 |
| BPNT1 | 5 |
| SLC6A6 | 5 |
| BRSK2 | 5 |
| AGAP2 | 5 |
| ABR | 5 |
| FBXO2 | 5 |

| | |
|---|---|
| HM13 | 5 |
| ARPC1A | 5 |
| ARL1 | 5 |
| ATXN10 | 5 |
| SIK3 | 5 |
| PRKCE | 5 |
| AIFM3 | 5 |
| CPD | 5 |
| MARCKSL1 | 5 |
| PURA | 5 |
| LYN | 5 |
| TMED1 | 5 |
| DNAJB4 | 5 |
| PSMA4 | 5 |
| PCBP1 | 5 |
| AK5 | 5 |
| TUBB3 | 5 |
| DBN1 | 5 |
| AHCY | 5 |
| ARPC2 | 5 |
| CHMP2A | 5 |
| NCKIPSD | 5 |
| CPNE6 | 5 |
| PSMD2 | 5 |
| ARHGEF2 | 5 |
| UBQLN2 | 5 |
| PPP2R2D | 5 |
| TMED5 | 5 |
| ITSN1 | 5 |
| RABGGTA | 5 |
| OGT | 5 |
| SLC12A2 | 5 |
| PSPH | 5 |
| PAICS | 5 |
| PPP2R5E | 5 |
| ARPC5L | 5 |
| PABPC1 | 5 |
| HUWE1 | 5 |
| NUDC | 5 |
| YWHAH | 5 |
| YWHAB | 5 |
| LGALSL | 5 |
| DLG1 | 5 |
| PGAM1 | 5 |
| DNAJC6 | 5 |
| PGP | 5 |
| EFHD2 | 5 |

| | |
|---|---|
| RAB1B | 5 |
| GABARAPL1 | 5 |
| CPNE4 | 5 |
| GNAL | 5 |
| RAB6B | 5 |
| DNAJA1 | 5 |
| CHGA | 5 |
| SHISA4 | 5 |
| PPM1H | 5 |
| HINT3 | 5 |
| RGS8 | 5 |
| ARL8A | 5 |
| APP | 5 |
| PCBP2 | 5 |
| TMX1 | 5 |
| TPPP3 | 5 |
| CLEC2L | 5 |
| CLCN3 | 5 |
| CNRIP1 | 5 |
| KCNK4 | 5 |
| DPYSL5 | 5 |
| CACNB3 | 5 |
| GNAQ | 5 |
| ATP1A3 | 5 |
| PLCB1 | 5 |
| GNAO1 | 5 |
| VCP | 5 |
| AP2B1 | 5 |
| PAFAH1B3 | 5 |
| PAFAH1B2 | 5 |
| GNAZ | 5 |
| SLC8A1 | 5 |
| DPP10 | 5 |
| INPP5A | 5 |
| GDA | 5 |
| GOT1 | 5 |
| ABCG2 | 5 |
| PRKACA | 5 |
| OLA1 | 5 |
| SIRT2 | 5 |
| SYT1 | 5 |
| ATP6V1B2 | 5 |
| FSCN1 | 5 |
| CDC42 | 5 |
| PSMA2 | 5 |
| PPP1R7 | 5 |
| MYH10 | 5 |

| | |
|---|---|
| PEBP1 | 5 |
| FKBP1A | 5 |
| PSMA6 | 5 |
| GRM2 | 5 |
| AARS1 | 5 |
| FRRS1L | 5 |
| CACNA2D3 | 5 |
| SLK | 5 |
| CAV1 | 5 |
| MDH1 | 5 |
| YWHAZ | 5 |
| PPIA | 5 |
| YWHAE | 5 |
| PRDX2 | 5 |
| PFN2 | 5 |
| YWHAG | 5 |
| APPL1 | 5 |
| PKM | 5 |
| KCNMA1 | 5 |
| SLC30A3 | 5 |
| PSMB2 | 5 |
| ACO1 | 5 |
| POMGNT2 | 5 |
| HNRNPK | 5 |
| GNB2 | 5 |
| MLEC | 5 |
| VPS29 | 5 |
| AP1G1 | 5 |
| EPB41L1 | 5 |
| RAB12 | 5 |
| RAB2A | 5 |
| ATP2B2 | 5 |
| WDR1 | 5 |
| TPI1 | 5 |
| GDI2 | 5 |
| SEPTIN11 | 5 |
| ARPC5 | 5 |
| RAB7A | 5 |
| ARPC4 | 5 |
| GDI1 | 5 |
| CAP2 | 5 |
| DTD1 | 5 |
| XPNPEP1 | 5 |
| RTN3 | 5 |
| PRMT8 | 5 |
| GPHN | 5 |
| CPNE2 | 5 |

| | |
|---|---|
| ERLIN2 | 5 |
| TOLLIP | 5 |
| LAMP5 | 5 |
| WDR48 | 5 |
| SLC3A2 | 5 |
| ATP6V0D1 | 5 |
| ATP6V0A1 | 5 |
| DNAJB11 | 5 |
| CAPZA2 | 5 |
| PCYOX1L | 5 |
| SEC11A | 5 |
| FKBP2 | 5 |
| RPN1 | 5 |
| DDOST | 5 |
| NCEH1 | 5 |
| SLC17A7 | 5 |
| TSPAN7 | 5 |
| OSTC | 5 |
| CA4 | 5 |
| THY1 | 5 |
| DPP4 | 5 |
| MAGI2 | 5 |
| IFNGR1 | 5 |
| ITGB1 | 5 |
| RALGAPB | 5 |
| POSTN | 5 |
| HPX | 5 |
| RASGRF1 | 5 |
| NUTF2 | 5 |
| BMPR2 | 5 |
| NTM | 5 |
| ERGIC1 | 5 |
| PSMB8 | 5 |
| RAB18 | 5 |
| HINT1 | 5 |
| PSMB5 | 5 |
| TCAIM | 5 |
| ARVCF | 5 |
| GOLT1B | 5 |
| APOO | 6 |
| TBRG4 | 6 |
| NDUFB5 | 6 |
| SAMM50 | 6 |
| BDH1 | 6 |
| AGK | 6 |
| ATP5F1C | 6 |
| ATP5F1E | 6 |

| | |
|---|---|
| ATP5PD | 6 |
| SSBP1 | 6 |
| ALDH5A1 | 6 |
| NDUFS4 | 6 |
| MPC1 | 6 |
| SFXN1 | 6 |
| SLC25A31 | 6 |
| OGDH | 6 |
| UQCRQ | 6 |
| MTCH1 | 6 |
| MAOA | 6 |
| UQCC1 | 6 |
| UQCRH | 6 |
| CKMT1 | 6 |
| TOMM40L | 6 |
| SDHB | 6 |
| CISD1 | 6 |
| COX7A2 | 6 |
| UQCRFS1 | 6 |
| NDUFS2 | 6 |
| UQCRC1 | 6 |
| GPD2 | 6 |
| PNPLA8 | 6 |
| ABCB8 | 6 |
| DNAJC11 | 6 |
| C1QB | 6 |
| MRPL21 | 6 |
| ABCB7 | 6 |
| EXOG | 6 |
| PDHA1 | 6 |
| GLUD1 | 6 |
| ABAT | 6 |
| OPA1 | 6 |
| SFXN5 | 6 |
| NIPSNAP3B | 6 |
| ATP5IF1 | 6 |
| IDH3G | 6 |
| NDUFV2 | 6 |
| NDUFA12 | 6 |
| NDUFA8 | 6 |
| NDUFS1 | 6 |
| NDUFS8 | 6 |
| NDUFS7 | 6 |
| UQCRC2 | 6 |
| MTCH2 | 6 |
| SDHA | 6 |
| NDUFA5 | 6 |

| | |
|---|---|
| PHB2 | 6 |
| COX6A1 | 6 |
| MICOS10 | 6 |
| DECR1 | 6 |
| MTND5 | 6 |
| OAT | 6 |
| IMMT | 6 |
| ATP5ME | 6 |
| SLC25A12 | 6 |
| ATP5F1B | 6 |
| ATP5F1A | 6 |
| SLC25A18 | 6 |
| SLC25A23 | 6 |
| COX5A | 6 |
| CYC1 | 6 |
| COX6C | 6 |
| MDH2 | 6 |
| MPC2 | 6 |
| NIPSNAP1 | 6 |
| ATP5PB | 6 |
| FAM162A | 6 |
| NDUFA4 | 6 |
| NDUFB3 | 6 |
| SLC25A3 | 6 |
| SLC25A11 | 6 |
| SLC25A4 | 6 |
| GOT2 | 6 |
| AFG3L2 | 6 |
| ATP5MK | 6 |
| NIPSNAP2 | 6 |
| ATP5MF | 6 |
| MGST3 | 6 |
| VDAC2 | 6 |
| VDAC3 | 6 |
| VDAC1 | 6 |
| FAM210A | 6 |
| NDUFA13 | 6 |
| COX7C | 6 |
| NDUFV1 | 6 |
| NDUFB2 | 6 |
| COX4I1 | 6 |
| NDUFB4 | 6 |
| COX5B | 6 |
| NDUFA9 | 6 |
| C1QBP | 6 |
| NDUFA10 | 6 |
| COX6B1 | 6 |

| | |
|---|---|
| PRDX3 | 6 |
| DLAT | 6 |
| NDUFA1 | 6 |
| NDUFB8 | 6 |
| NDUFA2 | 6 |
| CS | 6 |
| NDUFB10 | 6 |
| DLST | 6 |
| IDH3A | 6 |
| SOD2 | 6 |
| NDUFB6 | 6 |
| NDUFS3 | 6 |
| LETM1 | 6 |
| UQCRB | 6 |
| DLD | 6 |
| NDUFA6 | 6 |
| MFF | 6 |
| GHITM | 6 |
| MTND4 | 6 |
| NDUFA11 | 6 |
| MCU | 6 |
| NDUFB9 | 6 |
| SLC25A5 | 6 |
| NDUFB7 | 6 |
| ATP5MPL | 6 |
| NT5DC3 | 6 |
| BTBD17 | 6 |
| MT-CYB | 6 |
| MTND1 | 6 |
| MTCO1 | 6 |
| TMEM65 | 6 |
| NDUFS5 | 6 |
| NDUFS6 | 6 |
| MTCO2 | 6 |
| NDUFC2 | 6 |
| SLC25A22 | 6 |
| ETFDH | 6 |

#### References


1 D'Acunzo, P. *et al.* Isolation of mitochondria-derived mitovesicles and subpopulations of microvesicles and exosomes from brain tissues. *Nat Protoc* **17**, 2517-2549, doi:10.1038/s41596-022-00719-1 (2022).

2 Voorbraeck, L. *et al.* Astrocyte-neuron mitochondrial transfer via mitoEVs supports neuronal energy metabolism and is impaired in early Alzheimer's disease. *bioRxiv*, [Preprint], doi:10.64898/2026.03.09.710050.

3 Zhang, X. *et al.* Proteome-wide identification of ubiquitin interactions using UbIA-MS. *Nat Protoc* **13**, 530-550, doi:10.1038/nprot.2017.147 (2018).

4 Blondel, V., Guillaume, J., Lambiotte, R. & Lefebvre, E. Fast unfolding of communities in large networks. *Journal of statistical mechanics: theory and experiment* **10**, P10008 (2008).

5 Mijalkov, M. *et al.* BRAPH: A graph theory software for the analysis of brain connectivity. *PLoS One* **12**, e0178798, doi:10.1371/journal.pone.0178798 (2017).

6 Tiunn, I. *et al.* BRAPH 2: a flexible, open-source, reproducible, community-oriented, easy-to-use framework for network analyses in neurosciences. *SoftwareX* **35**, 102791 (2026).

7 Lancichinetti, A. & Fortunato, S. Consensus clustering in complex networks. *Sci Rep* **2**, 336, doi:10.1038/srep00336 (2012).

8 Royal Statistical Society (Great Britain). volumes (Blackwell Publishing Royal Statistical Society John Wiley & Sons, London, U.K. London, 1998).

9 Kanehisa, M. & Goto, S. KEGG: kyoto encyclopedia of genes and genomes. *Nucleic Acids Res* **28**, 27-30, doi:10.1093/nar/28.1.27 (2000).

10 Thomas, P. D. *et al.* PANTHER: Making genome-scale phylogenetics accessible to all. *Protein Sci* **31**, 8-22, doi:10.1002/pro.4218 (2022).

11 Ragueneau, E. *et al.* The Reactome Knowledgebase 2026. *Nucleic Acids Res* **54**, D673-D681, doi:10.1093/nar/gkaf1223 (2026).

12 Rath, S. *et al.* MitoCarta3.0: an updated mitochondrial proteome now with sub-organelle localization and pathway annotations. *Nucleic Acids Res* **49**, D1541-D1547, doi:10.1093/nar/gkaa1011 (2021).

13 UniProt, C. UniProt: the Universal Protein Knowledgebase in 2025. *Nucleic Acids Res* **53**, D609-D617, doi:10.1093/nar/gkae1010 (2025).

14 Hein, M. Y. *et al.* Global organelle profiling reveals subcellular localization and remodeling at proteome scale. *Cell* **188**, 1137-1155 e1120, doi:10.1016/j.cell.2024.11.028 (2025).

15 Thul, P. J. *et al.* A subcellular map of the human proteome. *Science* **356**, doi:10.1126/science.aal3321 (2017).

16 Karlsson, M. *et al.* A single-cell type transcriptomics map of human tissues. *Sci Adv* **7**, doi:10.1126/sciadv.abh2169 (2021).

17 Fowler, S. L. *et al.* Tau filaments are tethered within brain extracellular vesicles in Alzheimer's disease. *Nat Neurosci* **28**, 40-48, doi:10.1038/s41593-024-01801-5 (2025).

18 Montine, T. J. *et al.* National Institute on Aging-Alzheimer's Association guidelines for the neuropathologic assessment of Alzheimer's disease: a practical approach. *Acta Neuropathol* **123**, 1-11, doi:10.1007/s00401-011-0910-3 (2012).

19 Ferreira, I. L., Carmo, C., Naia, L., S, I. M. & Cristina Rego, A. Assessing Mitochondrial Function in In Vitro and Ex Vivo Models of Huntington's Disease. *Methods Mol Biol* **1780**, 415-442, doi:10.1007/978-1-4939-7825-0_19 (2018).
